# *Phytoene Synthase 2* Can Compensate for the Absence of *Psy1* in Pepper Fruit (*Capsicum annuum*)

**DOI:** 10.1101/797977

**Authors:** So-Jeong Jang, Hyo-Bong Jeong, Ayoung Jung, Min-Young Kang, Suna Kim, Sun-Hwa Ha, Jin-Kyung Kwon, Byoung-Cheorl Kang

**Affiliations:** Department of Plant Science, Plant Genomics & Breeding Institute, and Research Institute of Agriculture and Life Sciences, Seoul National University, Seoul 151-921, Republic of Korea; Food and Nutrition in Home Economics, Korea National Open University, 169 Dongsung-Dong, Jongno-Gu, Seoul 110-791, Republic of Korea; Department of Genetic Engineering and Graduate School of Biotechnology, College of Life Sciences, Kyung Hee University, Giheung-gu, Yongin-si, Gyeonggi-do 17104, Republic of Korea; Institutes of Green-bio Science and Technology, Seoul National University, Pyeongchang 232-916, Republic of Korea

**Keywords:** capsanthin-capsorubin synthase, carotenoid, color complementation, mature fruit color, pepper (*Capsicum* spp.), phytoene synthase, virus-induced gene silencing (VIGS).

## Abstract

*Phytoene synthase 1* (*PSY1*) and *Capsanthin-capsorubin synthase* (*CCS*) are two major genes responsible for fruit color variation in pepper (*Capsicum* spp.), although fruit colors cannot be explained by variations in these two genes alone. Furthermore, the role of *PSY2* in fruit color development in pepper is unknown. Here, we used a systemic approach to discover the genetic factors responsible for the yellow fruit color of *C. annuum* ‘MicroPep Yellow’ (MY) and to reveal the role of *PSY2* in fruit color. We detected a complete deletion of *PSY1* and a retrotransposon insertion in *CCS* in MY. Despite the loss of *PSY1* and *CCS* function, the MY and mutant F_2_ plants from a cross between MY and the MicroPep Red (MR) accumulated basal levels of carotenoids, indicating that other *PSY* genes may complement the loss of *PSY1*. A qRT-PCR analysis demonstrated that *PSY2* is constitutively expressed in both MR and MY fruits, and a color complementation assay using *Escherichia coli* revealed that PSY2 is capable of biosynthesizing a carotenoid. Virus-induced gene silencing of *PSY2* in MY resulted in white fruits. These findings suggest that PSY2 can compensate for the absence of PSY1 in fruit, resulting in the yellow color of MY fruits.

**Highlight:** We reveal the novel function of *PSY2* in the development of yellow pepper fruit coloration using a *psy1* knockout mutant. This gene function was not previously identified in solanaceous crops.

## Introduction

Peppers (*Capsicum* spp.) are one of the most popular fruits in the world. Among the traits of pepper fruits, color is one of the most important for its aesthetic value and nutritional benefits. The distinct colors of mature fruits result from the chloroplast-to-chromoplast conversion during fruit ripening. Since chromoplasts possess a higher capacity to biosynthesize and sequester carotenoids than chloroplasts, mature pepper fruits often display ivory to red colors depending on chromoplast development and carotenoid biosynthesis (Sun *et al*., 2018). Carotenoids belong to a group of tetraterpenoids derived from eight isoprene units, thus contain 40 carbons in their polyene backbone (Jackson *et al*., 2008). Carotenoids are essential for human nutrition and health as they provide a dietary source of provitamin A and serve as antioxidants to reduce the incidence of many diseases, including cardiovascular diseases and cancers (Fraser and Bramley, 2004). Carotenoids also play essential roles in plants, functioning in photosynthesis and photo-protection, and providing precursors for phytohormones such as abscisic acid and strigolactones (Nambara and Marion-Poll, 2005; Domonkos *et al*., 2013; Al-Babili and Bouwmeester, 2015).

Carotenoid biosynthesis begins with the formation of phytoene, catalyzed by PSY. There are at least two *PSY* genes in plants except *Arabidopsis thaliana*, which only has a single *PSY* gene. Multiple *PSY* genes in one plant can explain different carotenogenesis in various tissues; for example, in tomato (*Solanum lycopersicum*), *PSY1* functions in the fruits, *PSY2* in the leaves, and *PSY3* in the roots (Fantini *et al*., 2013, Wang *et al*., 2014; Yuan *et al*., 2015). Following a series of desaturation and isomerization steps, phytoene is converted into lycopene, which is the major carotenoid component in mature tomato fruits. By contrast, the red color of pepper fruits is caused by the accumulation of capsanthin and capsorubin, which derived from lycopene (Paran and van der Knaap, 2007). This pepper-specific process is regulated by CCS, which converts antheraxanthin and violaxanthin into capsanthin and capsorubin, respectively (Bouvier *et al*., 1994).

The three-locus model (*C1*, *C2*, and *Y*) was proposed to explain the color variation in pepper fruits (Hurtado-Hernandez and Smith, 1985). In this model, mature fruit colors can be classified into eight groups according to the allelic combinations of three genes: from white (*c1 c2 y*) to red (*C1 C2 Y*). An analysis of candidate carotenoid biosynthetic genes revealed that the *C2* and *Y* loci encode *PSY1* and *CCS*, respectively (Lefebvre *et al*., 1998; Popovsky and Paran 2000; Huh *et al*., 2001); however, the third locus, *C1*, is yet to be characterized. Since the identification of the genes encoded by the *C2* and *Y* loci, many studies have explored the color variation in pepper fruits based on the allelic variations of *PSY* and *CCS*. In general, peppers with yellow fruit have structural mutations in the coding region of *CCS* that lead to a premature stop codon (Lefebvre *et al*., 1998; Popovsky and Paran 2000; Ha *et al*., 2007; Li *et al*., 2013). The early translational termination of *CCS* can also result in orange fruit, as was observed in *C. annuum ‘*Fogo’ (Guzman *et al*., 2010). By contrast, Kim *et al*. (2010) revealed that a splicing mutation in *PSY1* impairs the activity of its enzyme product and leads to orange bonnet pepper fruits (*C. chinense* ‘Habanero’). Despite this, allelic variations of *PSY* and *CCS* cannot fully explain the differences in fruit color across all pepper accessions, as Jeong *et al*. (2018) previously described. Therefore, identification of the last color-determining gene, *C1*, is necessary for solving the riddle of fruit color variations in pepper.

To identify *C1* and validate the three-locus model for fruit color variation in pepper, studies have investigated the roles of other carotenoid biosynthetic genes. The VIGS analysis of capsanthin biosynthetic genes in detached red pepper fruits showed that, in addition to *PSY1* and *CCS*, *lycopene ß-cyclase* (*Lcyb)* and *ß-carotene hydroxylase (CrtZ-2)* were also required for the formation of the normal red color (Tian *et al*., 2014). In addition, Borovsky *et al*. (2013) reported that an ethyl methanesulfonate (EMS)-induced nucleotide substitution in *CrtZ-2* coding region led to the conversion of red into orange fruits in *C. annuum* ‘Maor’. Despite this, natural mutations of these genes have not been identified, therefore they are not candidates of the *C1* locus.

The carotenoid biosynthetic pathways and the functions of the genes involved have been elucidated using strains of *Escherichia coli* as heterologous hosts, in a so-called color complementation assay (Perry *et al*., 1986; Cunningham and Gantt, 2007). The pAC-BETA vector contains the *crtE*, *crtB*, *crtI*, and *crtY* carotenogenic genes from *Erwinia herbicola*, which can produce β-carotene when transformed into *E. coli* (Cunningham *et al*., 1996). By contrast, the pAC-85b vector does not contain *crtB*, a homolog of *PSY*, and when expressed in *E. coli*, β-carotene can only be produced when complemented with a gene encoding a PSY enzyme (Cunningham and Gantt, 2007). Through a complementation assay using pAC-85b, the function of the *PSY1* transcriptional variants of both barley (*Hordeum vulgar*e) and tomato has been revealed (Rodríguez-Suárez *et al*., 2011; Kang *et al*., 2014), and the catalytic activities of multiple PSY enzymes were also analyzed in loquat (*Eriobotrya japonica*) (Fu *et al*., 2014). Moreover, using a color complementation test using the pAC-LYC construct, which contain genes for lycopene production in papaya (*Carica papaya*), the role of *Lcyb* in fruit flesh color was confirmed (Blas *et al*., 2010). To date, a wider range of carotenoid-related enzymes, such as the carotenoid cleavage dioxygenases, have been analyzed using the above-mentioned approach (Kim *et al*., 2013). However, *PSY* genes of pepper have not been analyzed by color complementation assay yet.

In this study, seven carotenoid biosynthetic genes were analyzed to discover the genetic factor regulating yellow fruit color in *C. annuum* ‘MicroPep Yellow’ (MY). Among them, structural mutations in *PSY1* and *CCS* were identified in MY. Despite the associated functional knockout of *PSY1*, which catalyze the rate-limiting step of carotenoid biosynthesis, basal levels of carotenoids were still detected in MY using ultra-performance liquid chromatography (UPLC) analysis. We postulated that *PSY2* may contribute to the formation of the yellow color in MY fruits. To test this hypothesis, a qRT-PCR analysis of *PSY2* expression and a color complementation assay was conducted. The VIGS of *PSY2* was used to confirm the function of this gene in MY. The result showed that *PSY2* are expressed in all stage of fruit and functionally active gene. Here, we suggest that PSY2 can compensate the absence of PSY1 in pepper fruit.

## Materials and methods

### Plant materials

*C. annuum* accessions ‘MicroPep Red (MR)’ and ‘MicroPep Yellow (MY)’ were selected from the germplasm of Horticultural Crops Breeding and Genetics lab (Seoul National University, Republic of Korea). The mature fruits of MR and MY were red and yellow, respectively (**Fig. 1**). A total of 281 F_2_ individuals were obtained from a cross between MR and MY and grown at the greenhouse of Seoul National University (Suwon, Republic of Korea).

**Fig. 1.**
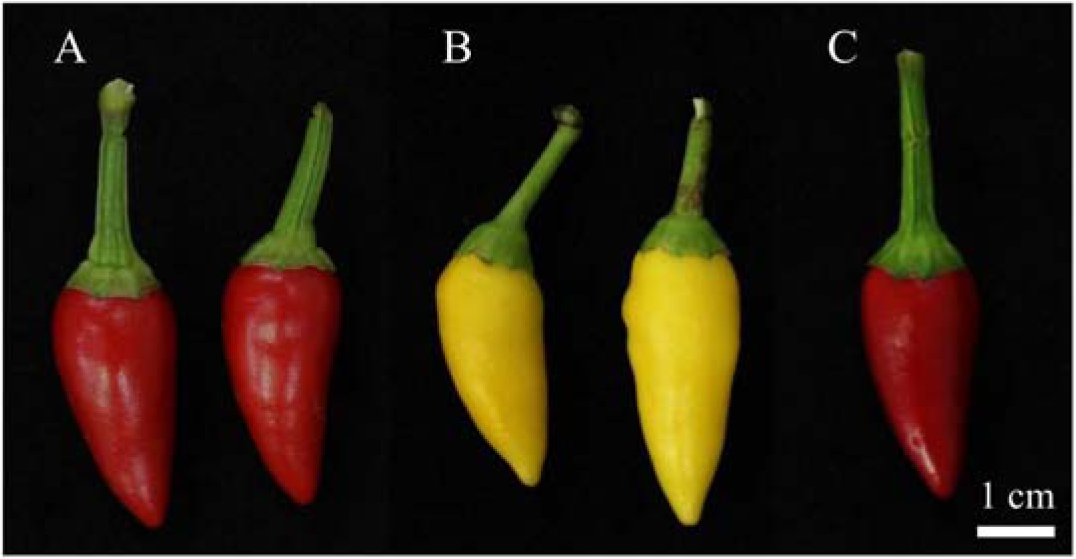
Mature pepper fruits of the genotypes used in this study. Mature fruit color of MicroPep Red (MR; A), MicroPep Yellow (MY; B), and an F_1_ hybrid derived from the MR × MY cross (C).

### Nucleic acid extraction

Genomic DNA (gDNA) was extracted from the young leaves of F_2_ individuals from a cross between MR and MY, and of the parental lines, using a modified cetyltrimethylammonium bromide (CTAB) method (Porebski *et al*., 1997). Leaf tissues were homogenized using 3-mm steel beads with TissueLyserII (Qiagen, Hilden, Germany). The gDNA concentration were measured and diluted to 20 ng/μL for further experiments.

Total RNA was extracted from target tissues, immediately frozen in liquid nitrogen and ground into a fine powder. The RNA was extracted using a MG RNAzol kit (MGmed, Seoul, Republic of Korea), following manufacturer’s instructions. The cDNA was synthesized from 2 μg RNA using an EasyScript Reverse Transcriptase kit (TransGen, Beijing, China) with oligo(dT) primers. The resulting cDNAs were used for the expression analysis.

### Candidate gene analysis

Reference sequences of seven carotenogenic genes in pepper were obtained from the Sol Genomics Network (https://solgenomics.net/). *PDS* was not included as its mutation results in distinct effect. The pepper *PSY2* and *PSY3* sequences have not been identified previously; therefore, the tomato *PSY2* and *PSY3* sequences were BLAST-searched against the pepper genome (CM334 v1.55). As a result, the *CA02g20350* (92% sequence identity with tomato *PSY2*; *Solyc02g081330*) and *CA01g12040* (93% sequence identity with tomato *PSY3*; *Solyc01g005940*) coding sequences were obtained. Primers were designed to amplify the full length of these candidate genes (**Table 1**), and a PCR was performed using PrimeSTAR GXL DNA polymerase (Takara Bio, Kusatsu, Japan). The resulting amplicons were separated on a 1% agarose gel and the DNA was recovered using a LaboPass PCR clean-up kit (Cosmo Genetech, Seoul, Republic of Korea). Sanger sequencing was performed at Macrogen (Seoul, Republic of Korea). The nucleotide sequences were analyzed using Lasergene’s SeqMan program.

**Table 1.**
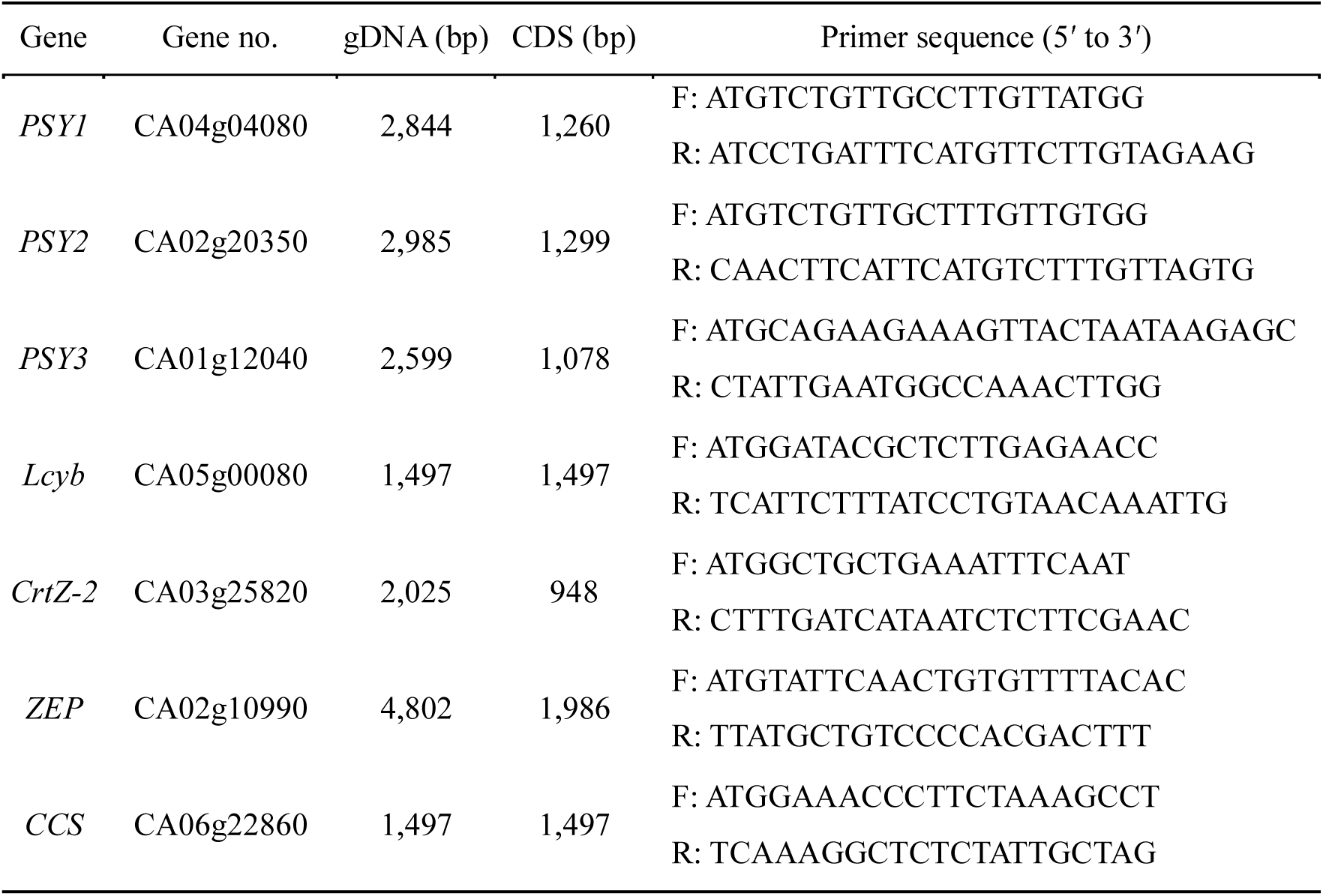
List of primers used to amplify the carotenoid biosynthetic genes.

### Genome walking from a *PSY1*-adjacent gene

To reveal the structural variation of *PSY1* in MY, genome walking was performed starting from *PSY1*-adjacent gene. Separate aliquots of MY gDNA were digested using four restriction enzymes: *Dra*I, *Eco*RV, *Pvu*II, and *Stu*I. Each digested DNA was separately ligated to a genome walker adaptor of Universal GenomeWalker 2.0 kit (Takara Bio). After the library construction, consecutive PCR amplifications were performed. For the primary PCR, an outer adaptor primer (AP1) and an outer gene-specific primer (GSP1) were used. The primary PCR product was diluted 50× and used as a template for the secondary PCR with a nested adaptor primer (AP2) and a nested gene-specific primer (GSP2). *Capana04g002520* is located in the adjacent 5′ region of *PSY1* (*Capana04g002519*) based on the Zunla reference genome (v2.0), so gene-specific primers were first designed to target the coding region of *Capana04g002520*. The secondary PCR amplicons were purified and sequenced by Macrogen. The next set of gene-specific primers was subsequently designed based on the sequenced genomic region. The primary PCR conditions were as follows: eight cycles of 94°C for 25 s and 72°C for 3 min, then 32 cycles of 94°C for 25 s and 67°C for 3 min. The secondary PCR conditions were as follows: six cycles of 94°C for 25 s and 72°C for 3 min, then 19 cycles of 94°C for 25 s and 67°C for 3 min.

### Long-range PCR amplification of *CCS*

To investigate whether an insertion was present in MY *CCS*, a long-range PCR was performed using the primers for CCS amplification (**Table 1**) and PrimeSTAR GXL DNA polymerase (Takara Bio). A two-step PCR was performed for 30 cycles of 98°C for 10 s and 68°C for 10 min. The resulting amplicons were purified and cloned into a modified T-blunt vector (SolGent, Daejeon, Republic of Korea). The plasmid DNA was extracted using an AccuPrep plasmid mini extraction kit (Bioneer, Daejeon, Republic of Korea) and sequenced by Macrogen.

### Sequence-characterized amplified region (SCAR) marker development for the structural variations of *PSY1* and *CCS*

SCAR markers for *PSY1* and *CCS* were designed based on the identified structural variations. A common (COM) primer and an allele-specific primer for the MR or MY genotypes were paired for gene amplification (**Table S1**). The SCAR markers were used to genotype the MY x MR F_2_ plants, and to screen germplasm accessions of interest which *PSY1* was not amplified in another study (Jeong *et al*., 2018). SCAR marker PCR condition was as follows: 35 cycles of 98°C for 10 s, 60°C for 30 s, and 72°C for 1 min. The resulting amplicons were separated on a 2% agarose gel.

### Expression analysis of *PSY2* using quantitative real-time PCR (qRT-PCR)

For the precise measurement of *PSY2* expression, a qRT-PCR was carried out using a LightCycler 480 system (Roche, Basel, Switzerland) using SYTO 9 stain (Thermo Fisher Scientific). The reaction mixture comprised a total volume of 20 μL, containing 2 μL 4× diluted cDNA, 0.3 μL R Taq (Takara Bio), 2 μL 10× PCR buffer, 2 μL 10 mM dNTPs, 0.5 μL 10 pmol primers, and 0.5 μL SYTO 9. The primer sequences were as follows: 5′-AGACAGAGGTGGAATTTTGGGTCT-3′ (forward) and 5′-CAAATTCCCCGGAAGCACA-3′ (reverse). The real-time PCR involved 45 cycles of 95°C for 10 s, 60°C for 20 s, and 72°C for 20 s. Three replicate reactions were performed. The *Actin* served as an internal control.

### Complementation assay for *PSY2* in *E. coli*

To analyze the enzymatic activity of PSY2, a complementation assay was conducted using pAC-85b (Addgene plasmid #53282). pAC-85b is derived from pAC-BETA (Addgene plasmid #53272), which contains *CrtE*, *CrtY*, *CrtI*, and *CrtB* (the carotenogenic gene cluster) from *Erwinia herbicola* for β-carotene production. pAC-85b lacks *PSY* homolog *CrtB* compared to pAC-BETA, meaning that *E. coli* transformed with pAC-85b can only produce β-carotene when complemented with a gene encoding a PSY. For the transgenic expression of *PSY1* and *PSY2*, pET-28a(+) (Novagen; Merck Millipore, Burlington, USA) was used (**Fig. S1**). To construct pET-PSY1 and pET-PSY2, the full-length cDNA fragments were amplified using the following primers with a 4-bp adapter and 6-bp restriction site (underlined): 5′-CACA<u>GGATCC</u>ATGTCTGTTGCCTTGTTATGG-3′ (*PSY1* forward) and 5′-CACA<u>GTCGAC</u>ATCCTGATTTCATGTTCTTGTAGAAG-3′ (*PSY1* reverse); 5′-CCAA<u>GGATCC</u>ATGTCTGTTGCTTTGTTGTGG-3′ (*PSY2* forward) and 5′-GGAA<u>GTCGAC</u>CAACTTCATTCATGTCTTTGTTAGTG-3′ (*PSY2* reverse). The amplicons were cloned into the *Bam*HI and *Sal*I sites of pET-28a(+). After the sequences were confirmed, pAC-85b and either pET-PSY1 or pET-PSY2 were co-transformed into the *E. coli* BL21(DE3) competent cells (TransGen), and named PC1 and EXP respectively. The control cells were co-transformed with pAC-85b and pET-28a(+) for negative control (NC), and with pAC-BETA for positive control (PC2) for a visual confirmation of the β-carotene content.

The transformed *E. coli* cells were grown overnight at 28°C in 2 mL of a Luria-Bertani (LB) liquid medium containing 50 mg/mL kanamycin and 35 mg/mL chloramphenicol and used to inoculate 10 mL LB medium containing the same antibiotics. The cells were grown at 28°C to OD_600_ 0.5–0.7, then 50 μM isopropylthio-β-galactoside (IPTG) was added and the cells were further incubated in darkness at 16°C for 24 h. Cell pellets were harvested by centrifugation and the colors of the pellets were analyzed.

### Immunoblot assay of the PSY proteins produced by the transgenic *E. coli*

NC, EXP, and PC1 *E. coli* were grown at 16°C for 24 h, after which 2 mL of each sample was transferred into 1.5-mL tubes, harvested by centrifugation, and frozen at –20°C. The frozen cells were resuspended in 800 μL distilled water and 200 μL 5× sample buffer, then incubated at 98°C for 5 min for denaturation. The proteins were separated using SDS-PAGE at 130 V for 1 h 30 min. After the electrophoresis, one repeat was cut from the gel and transferred to an Amersham Hybond™ -P membrane (GE Healthcare, Chicago, USA), using manufacturer’s instructions. The other sample replicate was stained with Coomassie brilliant blue solution.

The membrane was blocked using TBST buffer containing 5% BSA for 1 h at room temperature (RT). An antibody was raised against the middle region of the tomato PSY1 in mouse (anti-SlPSY1 (M)), as a primary antibody to bind to PSY2 for amino acid sequence similarity between *Sl*PSY1 and *Ca*PSY2. Anti-mouse-IgG-HRP was used as a secondary antibody. The anti-PSY1 (M) antibody (1 mg/μL) was diluted 5000:1 using TBST buffer containing 5% BSA. The primary antibodies were applied to the blocked membrane, and incubated at RT overnight. The membranes were washed twice by shaking and twice on the rocker for 10 min with TBST buffer. The secondary antibody (1 mg/μL) was diluted 5000:1 with TBST buffer, and applied to the membrane for 3 h on rocker speed 1. The membrane was washed as described above. Clarity™ Western ECL substrate (Bio-Rad) and Fusion FX (Vilber Lourmat, Collegien, France) were used for signal development and detection following the manufacturers’ instructions. The expected PSY2 sizes were calculated using a protein molecular weight calculator (www.sciencegateway.org/tools/proteinmw.htm).

### Carotenoid extraction and UPLC analysis

The carotenoids were extracted using methods specific to the materials (either fruit pericarp or *E. coli*). For the pericarp tissues, the carotenoid extraction and saponification were conducted as described by Kim *et al*. (2016). For the *E. coli* transformants, cell pellets were obtained using centrifugation, from which the carotenoids were extracted with 600 μL HPLC-grade acetone (Honeywell, Charlotte, USA), as described by López-Emparán *et al*. (2014).

For the extracts derived from both the pericarps and *E. coli* cell pellets, the carotenoids were analyzed using an Acquity UPLC-H-Class system (Waters, Milford, MA, USA). The compounds were separated using an Acquity UPLC HSS T3 column (2.1 × 100, 1.8 μm) at 35°C. The mobile phase was a binary solvent system consisting of phase A (acetonitrile/methanol/methylene chloride, 65/25/10, v/v/v) and phase B (distilled water). The gradients were programmed according to the previously described method (Kim *et al*., 2016). For the qualitative and quantitative analyses, 11 carotenoid standards were purchased from Sigma-Aldrich (St. Louis, USA); antheraxanthin, capsanthin, capsorubin, lutein, neoxanthin, violaxanthin, zeaxanthin, α-carotene, α-cryptoxanthin, β-carotene, and β-cryptoxanthin.

### VIGS of *PSY2* in MY

Ligation independent cloning (LIC) was conducted for the construction of pTRV2-PSY2 as described by Kim *et al*. (2017). The partial coding sequence of *PSY2* was amplified with the following primers containing 15-bp adapter sequences (underlined): 5′-<u>CGACGACAAGACCCT</u>TGTTGCTTTGTTGTGGGTTG-3′ (forward) and 5′-<u>GAGGAGAAGAGCCCT</u>TCCCAAGTCCGAATATCTCAA-3′ (reverse). The purified PCR product and *Pst*I-digested pTRV2-LIC vector were treated with T4 DNA polymerase (Enzymatics, Beverly, USA) and 5× blue buffer in common, and 10 mM dATP or dTTP respectively. Both mixtures were incubated at 22°C for 30 min, then 75°C for 20 min. A total of 30 ng PCR product and 400 ng TRV2-LIC vector were mixed and incubated at RT for 15 min. The mixture was transformed into *E. coli* DH5α competent cells (TransGen). Plasmids were extracted, sequenced by Macrogen, and introduced into *Agrobacterium tumefaciens* strain GV3101 through electroporation at 2.0 kV. For the negative VIGS control, pTRV2-GFP was used. Both pTRV2-LIC and pTRV2-GFP vectors were kindly provided by Dr. Doil Choi, Seoul National University.

*Agrobacterium* carrying pTRV1, pTRV2-GFP, or pTRV2-PSY2 were grown overnight at 30°C in LB medium with 50 μg/mL rifampicin and kanamycin, then harvested using centrifugation. The cells were resuspended in 10 mM MES, 10 mM MgCl_2_, and 200 μM acetosyringone to a final OD_600_ of 0.7–0.8, and incubated in a rocking incubator at RT for 3 h. Cell cultures containing pTRV1 and either of the pTRV2 constructs were mixed in a 1:1 ratio and infiltrated into the abaxial side of both pepper cotyledons. The inoculated plants were incubated at 16°C in the dark for one day, then grown at 25°C under a16-h/8-h light/dark photoperiod.

### Expression analysis of *PSY2* in VIGS-treated fruit using RT-PCR

Mature fruits were harvested from pTRV2-GFP- or pTRV2-PSY2-inoculated MY plants at four months after inoculation. Segments of the mature pericarp displaying lighter colors were excised and their RNA was extracted. An RT-PCR was conducted to verify the gene silencing using the following reaction mixture: 2 μL 4× diluted cDNA, 0.3 μL EX Taq (Takara, Japan), 2.5 μL 10× PCR buffer, 2 μL 10 mM dNTPs, 0.5 μL 10 pmol primers, and 17.2 μL triple distilled water. The same primers used to amplify the entire *PSY2* gene were used (**Table 1**). The RT-PCR conditions were 28 cycles of 95°C for 30 s, 58°C for 30 s, and 72°C for 90 s. Three biological replicates and the *Actin* were used for reproducibility and internal control, respectively.

### Carotenoid extraction and saponification of VIGS-treated fruits

Carotenoids were extracted as described by Yoo *et al*. (2017), with some modifications and under dim light. The samples were put on ice to prevent carotenoid oxidation and degradation. The lighter-colored pTRV2-PSY2-inoculated pericarps and the pTRV2-GFP-inoculated pericarps were diced and freeze-dried. Pieces from two TRV2-PSY2-inoculated fruits were pooled together to obtain enough tissue for carotenoid extraction.

The freeze-dried tissues were ground, and about 50 mg of tissue powder was placed in a 2-mL tube with two glass beads. For the saponification, 600 μL tetrahydrofuran (THF), 375 μL petroleum ether, and 150 μL 25% NaCl was added to the samples, vortexed, and centrifuged for 3 min at 1,500 g at 4°C. The upper phase was transferred into a new 2-mL tube. A 500-μL aliquot of petroleum ether was added to the remaining sample, which were then centrifuged, and this second upper phase was combined with the first upper phase. The samples were then concentrated using vacuum concentrator. The dried samples were dissolved with 500 μL HPLC-grade acetone (Honeywell) using sonication and filtered using a 0.2-μm syringe filter (Acrodisc LC 13 mm syringe filter, PVDF membrane; Pall, NY, USA) and bottled in a 1.5-mL HPLC amber vial. The HPLC analysis of the carotenoids was performed by the NICEM chromatography lab (Seoul National University). Capsanthin, capsorubin, lutein, zeaxanthin, α-carotene, β-carotene, and β-cryptoxanthin were used as standards.

## Results

### MY harbors structural variations in *PSY1* and *CCS*

Two *C. annuum* accessions, MR and MY, were used to identify the genetic factors controlling the formation of yellow coloration in the mature pepper fruit. The seven carotenoid biosynthetic genes *PSY1*, *PSY2*, *PSY3*, *Lcyb*, *CrtZ-2*, *ZEP*, and *CCS*, were amplified from the gDNA using PCR to identify any differences between the two genotypes. The expected sizes of the amplicons are shown in **Table 1**. In both MR and MY, the expected amplicon sizes were obtained for *PSY2*, *PSY3*, *Lcyb*, *CrtZ-2*, and *ZEP*, and no sequence variations were discovered in these genes between the accessions. By contrast, no amplicons were obtained from MY for *PSY1* and *CCS* (**Fig. 2A**). These results indicated the possible existence of structural variations in the *PSY1* and *CCS* genes in MY.

**Fig. 2.**
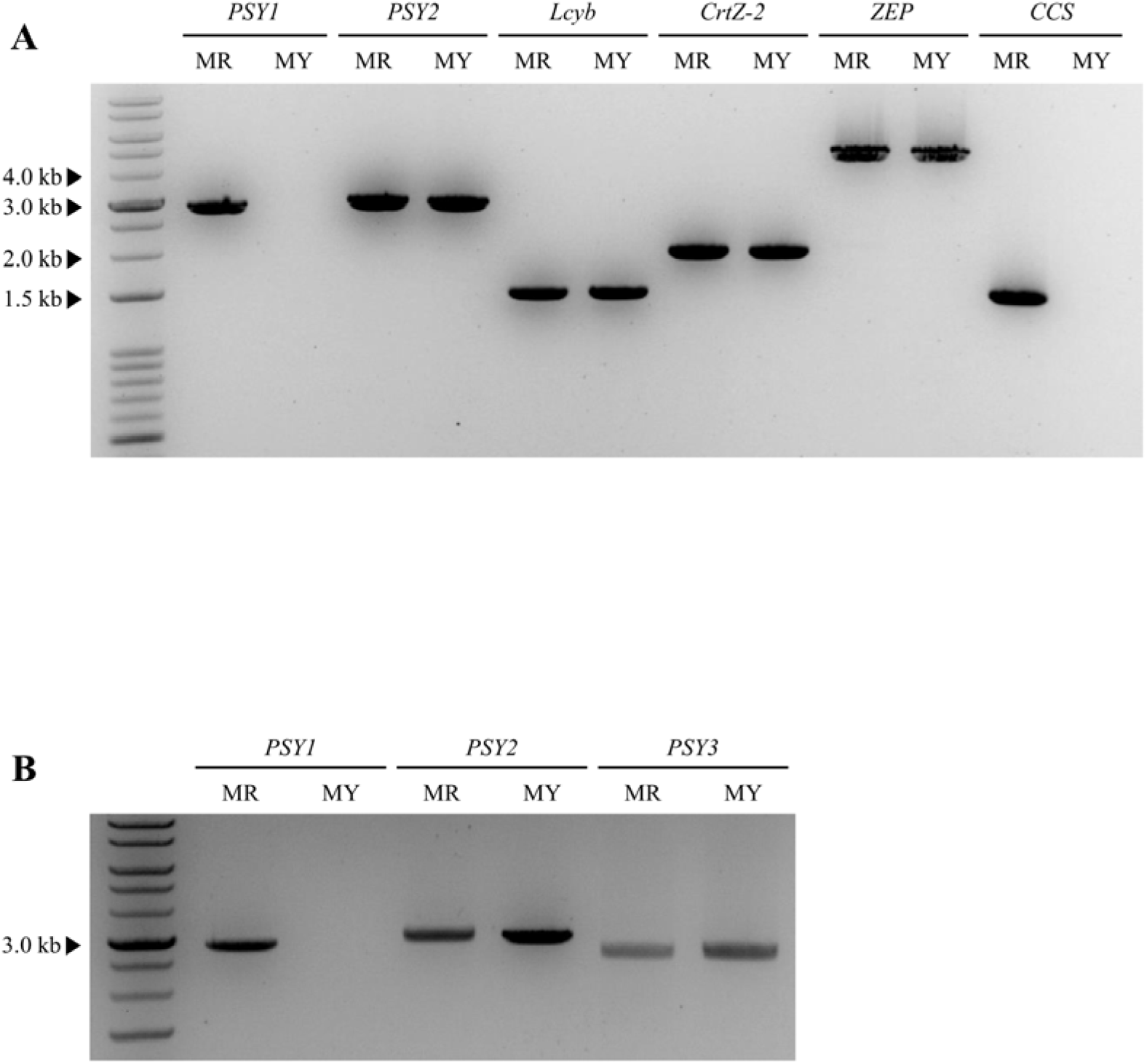
Polymorphism survey of the carotenoid biosynthetic genes. (A) gDNA amplification of six carotenoid biosynthetic genes using PCR. Primers were design to amplify the full length of each gene. From the two genotypes, the expected amplicon sizes were obtained for *PSY2, Lcyb, CrtZ-2,* and *ZEP*. No amplicon was obtained for *PSY1* and *CCS* in MY. (B) gDNA PCR amplification of the *PSY* genes. A difference in the amplification pattern between MY and MR was only detected for *PSY1*, for which no amplicon was obtained from MY.

### MY has a complete deletion of *PSY1* and a transposon insertion in *CCS*

To identify the structural variation of *PSY1* in MY, various regions of the gene, including the upstream and downstream sequences, were amplified using 12 sets of primers. No fragments of the expected size were obtained (data not shown), indicating the existence of a huge insertion or a deletion in the gene. To reveal the structural variation in *PSY1* in MY, genome walking was performed. Since target-specific primers are required for genome walking, the first primer set was designed using the gene *Capana04g0025200*, which is adjacent to *PSY1*. Using several rounds of genome walking, a 19,948-bp deletion including the entire *PSY1* genomic region was identified in the MY genome (**Fig. 3A**). To test commonality of this mutation among *C annuum* germplasm, we genotyped the 18 accessions showing yellow fruit color (Jeong *et al*., 2019) using the SCAR marker (**Table S1**) and showed that 15 of them had the same structural mutation as MY (**Fig. S4**).

**Fig. 3.**
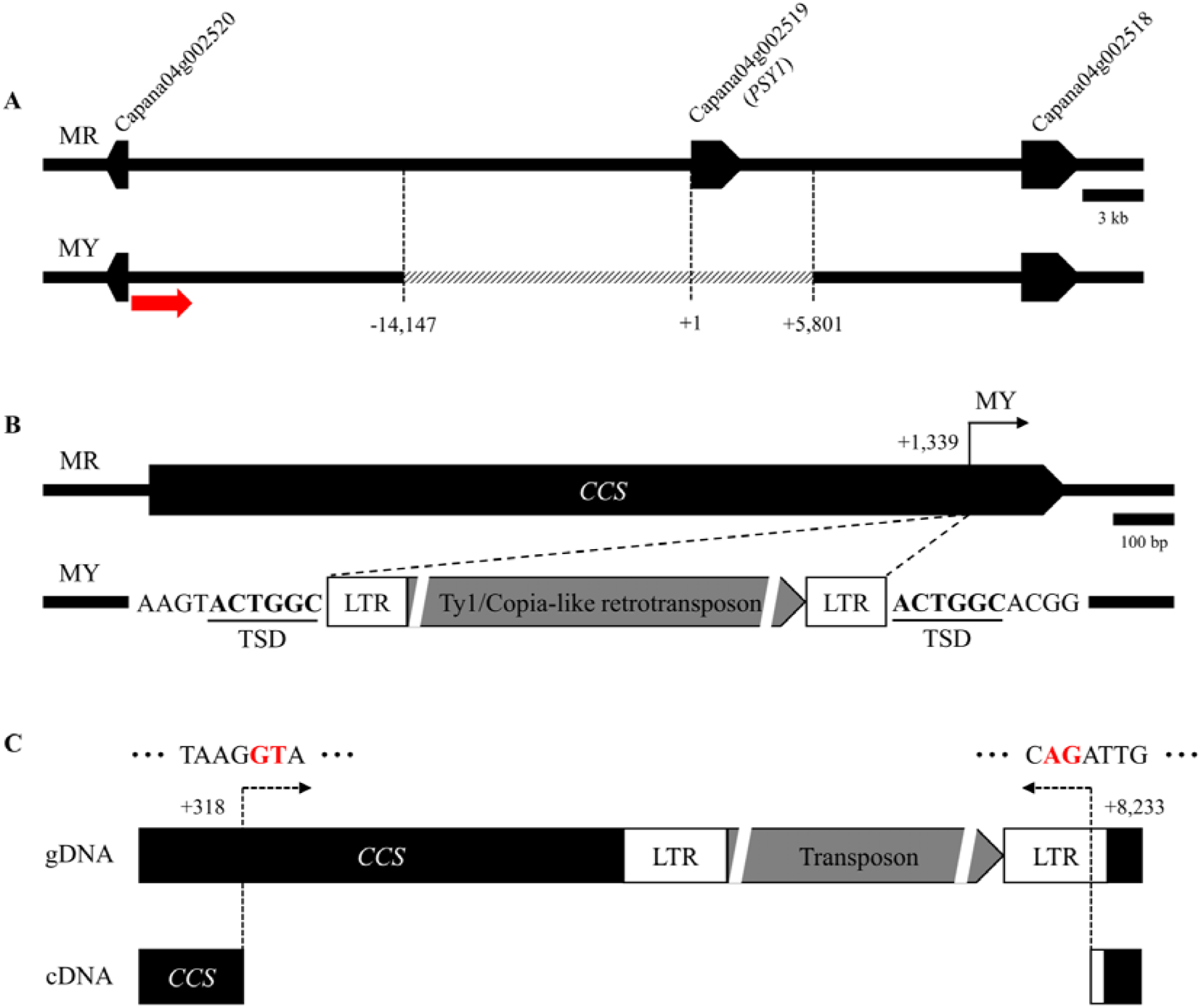
Structural variations of *PSY1* and *CCS* in MY. (A) An almost 20-kb deletion spanning the *PSY1* genomic region, revealed using genome walking from *Capana04g002520*. The deleted region is indicated with the diagonal lines. The primer used for the first genome walking run is indicated with a red arrow. (B) An almost 7-kb Ty1/Copia-like retrotransposon insertion was discovered 1,339 nt along *CCS* in MY. LTR, long terminal repeat. TSD, target site duplication. **(C)** Transcriptional variant of *CCS* in MY. Compared with the full *CCS* sequence in MR, much smaller bands were amplified from MY using RT-PCR. Such transcriptional variation was regarded as being caused by the provision of a conserved sequence in the splicing junction by the inserted sequence. The red letters represent the conserved sequences of the exon-intron junction. The genomic region indicated with a solid line denotes the region which is spliced out during transcription.

To investigate the structural variation of *CCS* in MY, a long-range PCR was conducted. Several regions of *CCS* were amplified normally (data not shown), indicating the presence of a large insertion that cannot be detected using regular PCR conditions. To overcome this limitation, the extension time of the PCR cycle was increased to 10 min, resulting in the amplification of a sequence containing a 6,750-bp insertion located 1,339 nt along *CCS* in MY (**Fig. 3B**). When this amplicon was sequenced and analyzed, the inserted sequence was observed to contain a Ty1/Copia-like retrotransposon, almost 1-kb long-terminal repeat (LTR) sequences, and a 6-bp target site duplication sequence (TSD)(ACTGGC) (**Fig. 3B**).

### MY produces a transcriptional variant of *CCS*

As MY contains the full *CCS* sequence, although interrupted by the transposon, an RT-PCR was performed to test whether functional transcripts of *CCS* were present in these plants. In MR, a 1,497-bp amplicon was obtained. By contrast, a much smaller amplicon (513 bp) was obtained from MY. A sequence analysis of this transcript showed that most of the transposon, along with 1 kb of *CCS*, were spliced out in MY (**Fig. 3C**). Such transcriptional variation might result from the provision of a conserved sequence in the splicing junction (5′-GT/AG-3′) by the inserted transposon sequence (Brown, 1986), which might be functioned as an active splicing junction of *CCS*. The resulting transcripts are expected to encode a truncated protein containing 115 amino acids, whereas functional CCS comprises 498 amino acids.

### Mutations in *PSY1* and *CCS* genes control fruit color

To test whether the structural variations of *PSY1* and *CCS* could result in the yellow fruit color of MY, an inheritance study was performed using a segregating population derived from a cross between MR and MY. All fruits from the F_1_ progeny were red, indicating that the red color is dominant over yellow. In the F_2_ population, red, orange, and yellow fruits showed a segregation ratio of 156:111:14 (**Table 2**). This ratio was consistent with an expected ratio of 9:6:1 (χ^2^ = 1.26, *P* = 0.59), suggesting that two genes epistatically interact to determine fruit color in this population.

**Table 2.** Segregation of phenotypes in the MR × MY F_2_ population.

| Population | Total<br>plants | Mature fruit<br>color | | | Expected ratio | $\chi^2$ |
| --- | --- | --- | --- | --- | --- | --- |
|  |  | Red | Orange | Yellow |  |  |
| MR × MY F<sub>2</sub> | 281 | 156 | 111 | 14 | 9:6:1 | 1.26 |

To test whether *PSY1* and *CCS* are involved in the determination of pepper fruit color, SCAR markers were used to genotype the F_2_ individuals (**Fig. S2**). Plants containing the wild-type alleles (*PSY1*/- and *CCS*/-) of both genes produced red fruits. By contrast, yellow fruits were produced by plants containing homozygous recessive alleles (*psy1*/*psy1 ccs*/*ccs*) for both genes. In plants containing either the *PSY1/- ccs/ccs* or *psy1/psy1 CCS/-* genotypes, the fruits were orange in color (**Table 3**).

**Table 3.** Co-segregation of genotypes and phenotypes in the MR × MY F_2_ population.

| Population | Size | Red | Orange | Orange | Yellow | Expected ratio | $\chi^2$ |
| --- | --- | --- | --- | --- | --- | --- | --- |
|  |  | *PSY1*/- | *PSY1*/- | *psy1/psy1* | *psy1/psy1* |  |  |
|  |  | *CCS*/- | *ccs/ccs* | *CCS*/- | *ccs/ccs* |  |  |
| MR × MY F<sub>2</sub> | 281 | 156 | 48 | 63 | 14 | 9:3:3:1 | 3.27 |

### Orange pepper fruits have either *PSY1* or *CCS* mutations

No phenotypic differences were observed between the orange-colored fruits produced by plants with different genotypes (*PSY1/- ccs/ccs* or *psy1*/*psy1 CCS*/-). Their carotenoid profiles were analyzed using UPLC. For each genotypic group, UPLC was performed with three biological replicates. The major carotenoid component of plants with the *PSY1*/- *CCS*/- genotype was capsanthin, whereas the main carotenoid in the plants harboring *psy1/psy1 ccs/ccs* were neoxanthin (**Fig. 4**). Two distinct differences were identified between the orange-colored fruits produced by plants containing the different genotypes. First, the total carotenoid contents of the fruits of the *PSY1/- ccs/ccs* genotype were four times higher than those of the fruits of the *psy1*/*psy1 CCS*/- genotype. Second, both capsanthin and capsorubin, the typical red pigments present in red peppers, were almost undetectable (0.35 mg/100 g dry weight (DW)) in the fruits of the *PSY1/- ccs/ccs* genotype, whereas these pigments accounted for almost 70% of total carotenoid levels in the *psy1/psy1 CCS/-* orange fruits (5.67 mg/100 g DW). These results indicated that, although the visible colors were similar between the two orange genotypes, the composition of pigments producing the orange color were different.

**Fig. 4.**
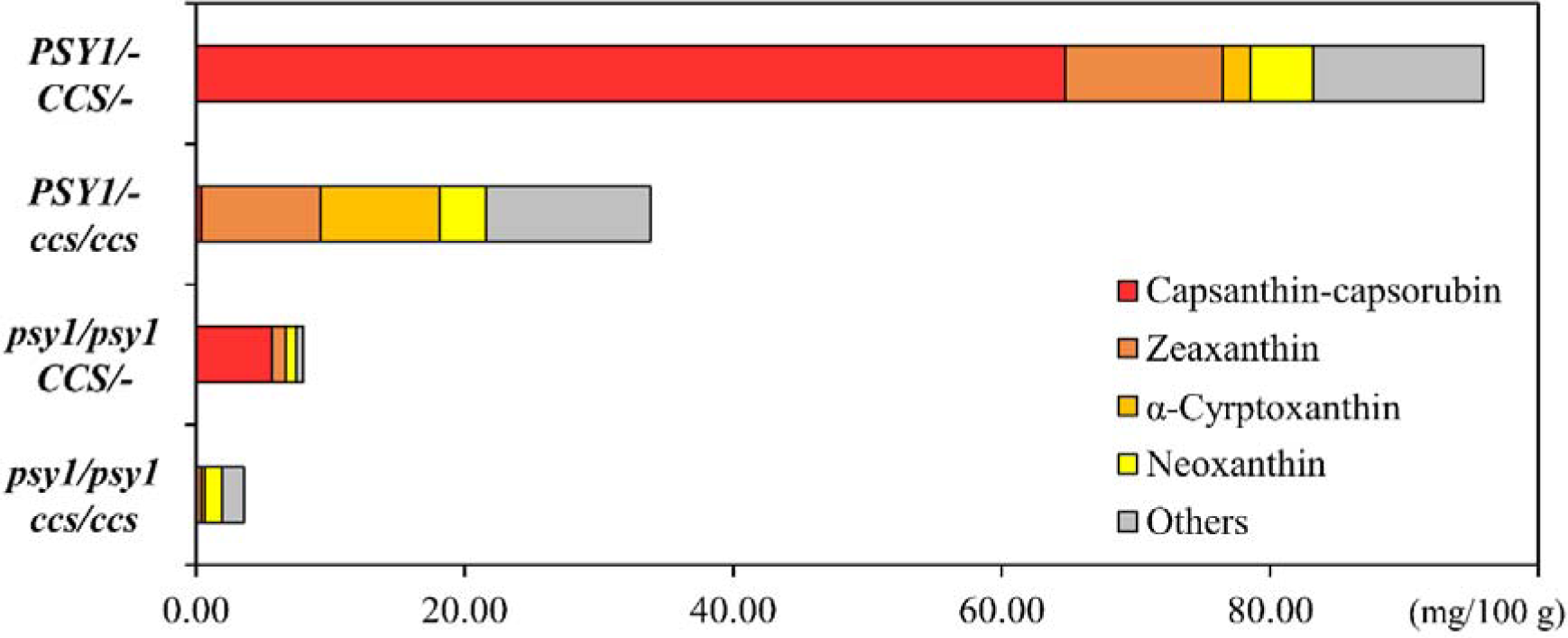
Carotenoid profiles of each F_2_ genotype. Among the 11 investigated carotenoid components, only the major carotenoids are described separately (capsanthin-capsorubin, zeaxanthin, α-cryptoxanthin, and neoxnathin), with the remaining minor carotenoids grouped together.

In the *psy1* knockout (*psy1*/*psy1*) plants, the carotenoids accumulated in the fruits regardless of the *CCS* genotype, although the total carotenoid content was about 10-fold lower than that of the plants containing functional a *PSY1* gene. In contrast to the *PSY1/- ccs/ccs* plants, the content of each carotenoid component was decreased in the *psy1/psy1 ccs/ccs* genotype. A similar trend was also observed in the *psy1/psy1 CCS/-* plants compared with the *PSY1/- CCS/-* plants, and the levels of most carotenoids except antheraxanthin were decreased. In conclusion, despite the impaired PSY1 activity, the carotenoid composition of the *psy1/psy1* plants was similar to the wild-type plants, although their levels were reduced (**Fig. 4**). This implies that other genes, such as the *PSY* homologs, might supplement the function of *PSY1*, regardless of *CCS*.

### *PSY2* is expressed in pepper fruits

Based on the UPLC data, it was postulated that *PSY2* or *PSY3* might be involved in carotenoid biosynthesis in the pepper fruits. To test this hypothesis, the expression levels of the *PSY* homologs were analyzed in the mature fruit pericarps using RT-PCR. The *PSY1* transcripts were only detected in MR. *PSY2* transcripts of the expected size were obtained from both MR and MY, while *PSY3* transcripts were not detected in either accession (**Fig. 5A**). These results demonstrate that *PSY2* is expressed in pepper fruits and may be involved in color formation. A further analysis using qRT-PCR confirmed that *PSY2* is expressed in the pericarp tissues, although the highest levels of expression were detected in the leaf tissues (**Fig. 5B**). The *PSY2* expression levels in the leaves and fruits were significantly higher in MY than in MR, indicating that *PSY2* may compensate for the loss of *PSY1* function in MY.

**Fig. 5.**
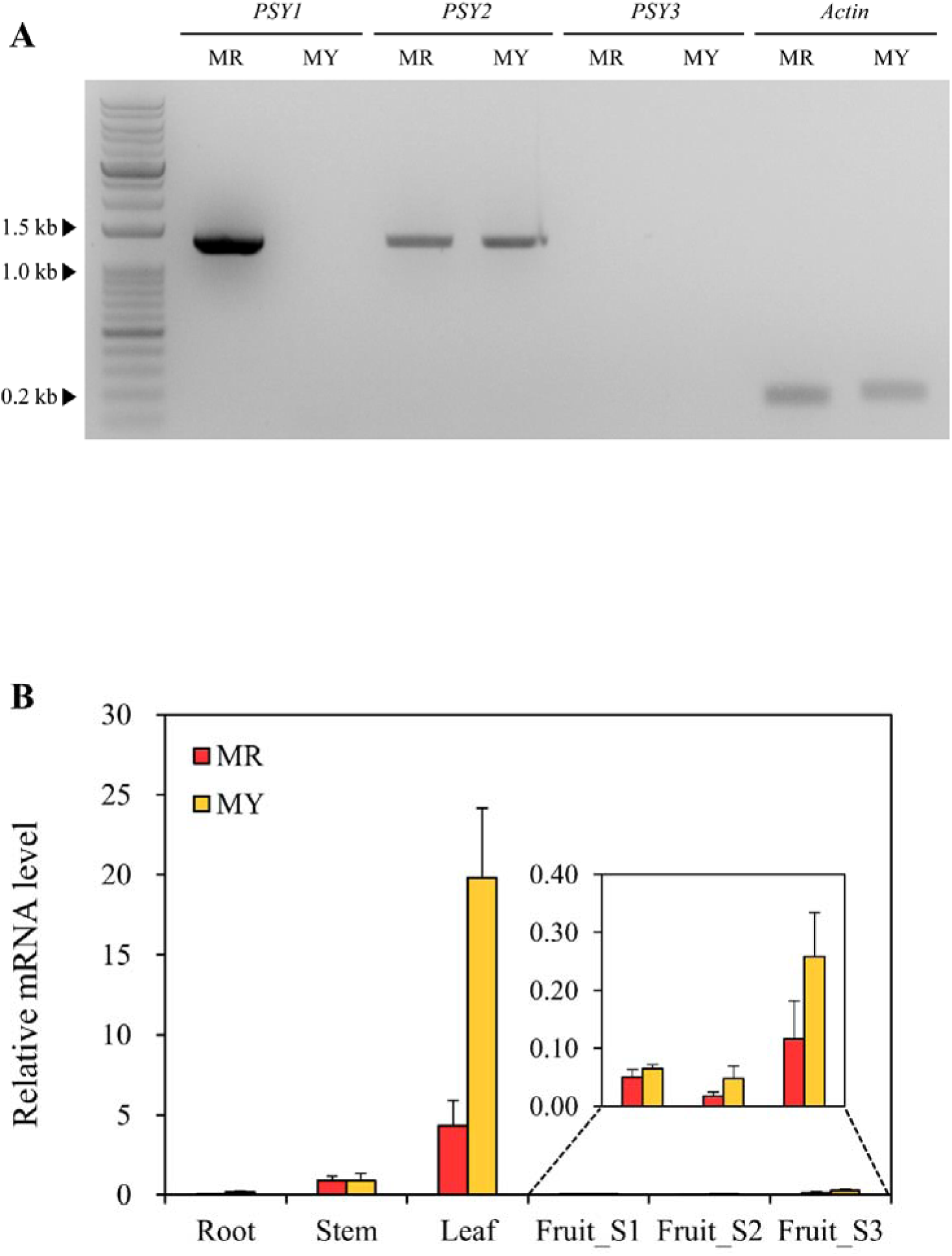
Expression pattern of the *PSY* genes in various tissues. (A) Expression analyses of the three *PSY* genes in the mature fruit pericarps. *Actin* serves as a control. (B) Expression analysis of *PSY2* in various tissues. Relative transcript levels of *PSY2* were determined using qRT-PCR, and normalized to the expression of *Actin*. The data are the means ± SD of three independent experiments. S, fruit developmental stage.

### PSY2 has the same enzymatic function as PSY1

The enzymatic function of PSY2 was tested using a complementation assay in *E. coli* BL21(DE3) cells carrying pAC-85b and pET-PSY2 (EXP). The same *E. coli* strain containing pAC-85b and pET-PSY1 was used as a positive control (PC1), while cells containing pAC-85b and the empty pET-28a(+) vector were used as the negative control (NC). For the comparison of the colored pigments produced by these cells, transformants producing β-carotene (pAC-BETA) were also used (**Fig. S1**). If PSY2 is able to biosynthesize the carotenoid backbone, the *E. coli* pellets would be orange in color due to the resulting β-carotene content. As shown in **Fig. 6A**, the *E. coli* pellets from EXP were orange, as were those of the two positive control groups, PC1 and PC2. UPLC analysis of the *E. coli* extracts was conducted to validate that the orange color of the experimental cells expressing *PSY2* was due to the accumulation of β-carotene. A clear β-carotene peak was detected in all cells except the negative control group (**Fig. 6B**). The β-carotene contents were 1.57 mg/100 g and 2.23 mg/100 g fresh weight (FW) in the cells containing pET-PSY2 and pET-PSY1, respectively. These results demonstrate that PSY2 has a similar catalytic activity to PSY1 in carotenoid biosynthesis.

**Fig. 6.**
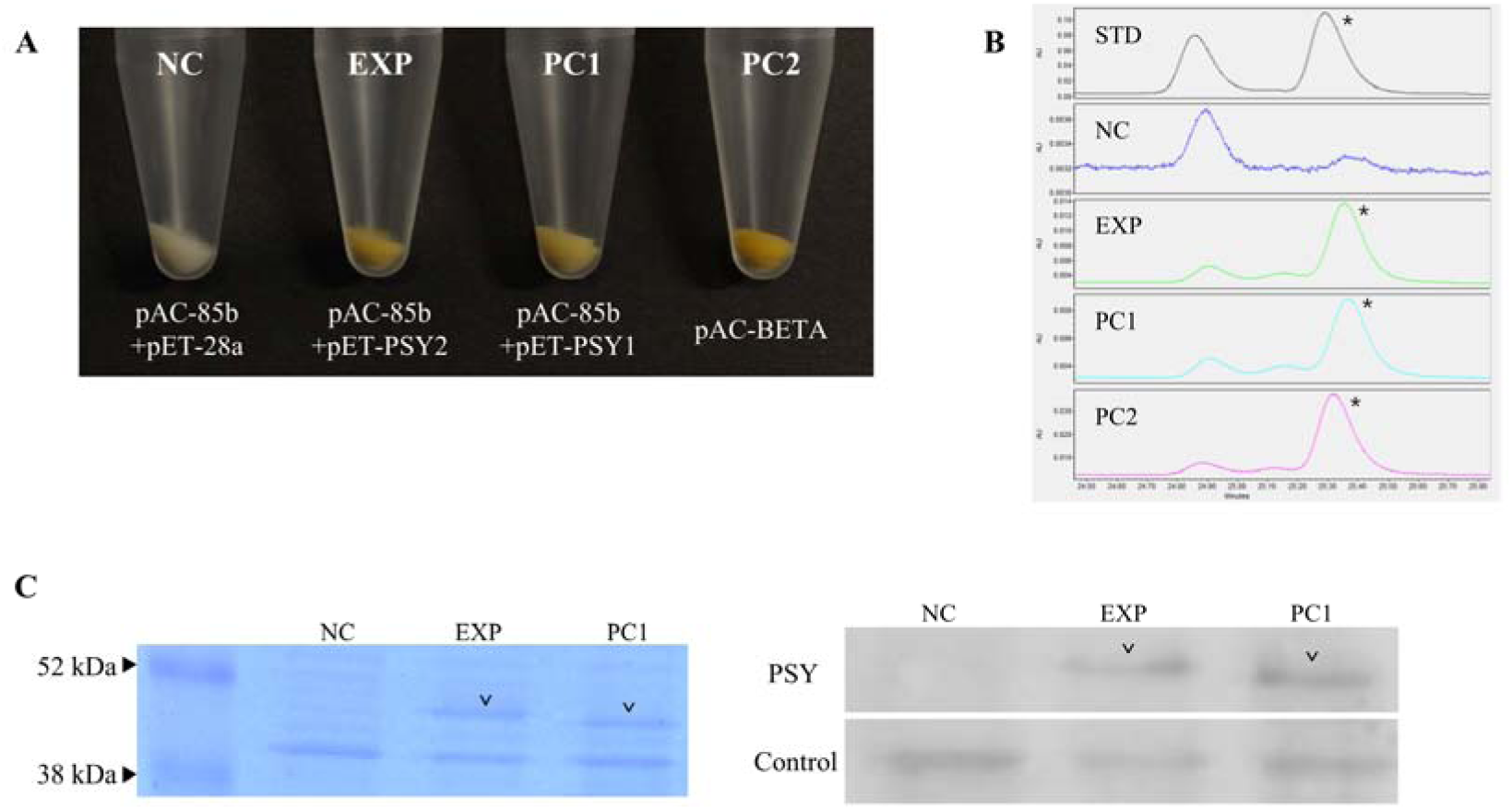
Complementation assay for *PSY2* in *E. coli*. (A) Expression of *PSY2* or *PSY1* in *E. coli* strains carrying pAC-85b. An *E. coli* strain carrying the empty vector alone served as a negative control. A β-carotene-accumulating *E. coli* strain served as a positive control for the color formation. (B) UPLC analysis of carotenoids extracted from each experimental group. The β-carotene peak is indicated with an asterisk. Au, absorbance units. STD, standard mixture of α- and β-carotenes. (C) Western blot analysis of *PSY* expression in transformed *E. coli*. Coomassie-stained SDS-PAGE gel (left) and the Western blot analysis (right) of the negative control (NC), the experimental group (EXP), and the positive control (PC1) using the anti-PSY-antibodies. The EXP and PC1-specific bands are indicated by arrowheads.

To validate the production of the PSY proteins in the *E. coli* cells, an immunoblot assay was conducted. A 2-mL aliquot of each cell culture was harvested, treated with SDS sample buffer, and ran on a 12% SDS-PAGE gel. One of the two replicated gels were stained with Coomassie blue. The EXP with pET-PSY2 and PC1 with pET-PSY1 specific band sizes were between 38 kDa and 52 kDa, as indicated in protein marker (**Fig. 6C**). The other sample replicate was transferred onto a nitrocellulose membrane and treated with an antibody against tomato *Sl*PSY1 (**Fig. S3**), which showed a high amino acid sequence similarity with *Ca*PSY1 (90.66%) and *Ca*PSY2 (82.96%; data not shown). Specific bands of around 48 kDa were detected in both EXP and PC1. The EXP band were slightly bigger than that of PC1, consistent with the expected sizes of CaPSY2 and *Ca*PSY1, 48.44 kDa and 47.07 kDa, respectively (**Fig. 6C**). These results demonstrate that the recombinant proteins are expressed in *E. coli*.

### Silencing of *PSY2* leads to lighter yellow fruit color

The function of *PSY2* was evaluated in MY plants using TRV-mediated VIGS, while *Agrobacterium* cells carrying pTRV2-GFP were used as a negative control. The symptoms of *PSY2* silencing were visualized at 15 days after inoculation (DAI). The pTRV2-GFP-inoculated plants did not display any significant differences to the wild type. The *PSY2*-silenced plants had a mosaic phenotype of pale and green patches on the leaves, due to the characteristics of the TRV vector (**Fig. 7A**). The flowers and fruits developed at about 60 and 75 DAI, respectively. There were no visible changes in the flower colors of both groups, but a slightly lighter peduncle color was observed in the TRV2-PSY2-inoculated plants (data not shown). In the pTRV2-GFP-inoculated plants, the immature fruits were a dark green color, whereas the immature fruits in the *PSY2-*silenced plants were white to pale green. The mature fruit color also showed a similar trend, as the pTRV2-GFP inoculated fruits showed an intense yellow color, whereas the *PSY2-*silenced fruits were white to ivory in color (**Fig. 7B**). RT-PCR was conducted to compare the expression levels of *PSY2* in the mature fruits of the pTRV2-GFP-inoculated and *PSY2*-silenced plants. While the fruits of the pTRV2-GFP-inoculated plants expressed *PSY2,* there was almost no detectable *PSY2* expression in the *PSY2-*silenced fruit tissues, indicating the successful post-transcriptional gene silencing (PTGS) of the target gene (**Fig. 7C**).

**Fig. 7.**
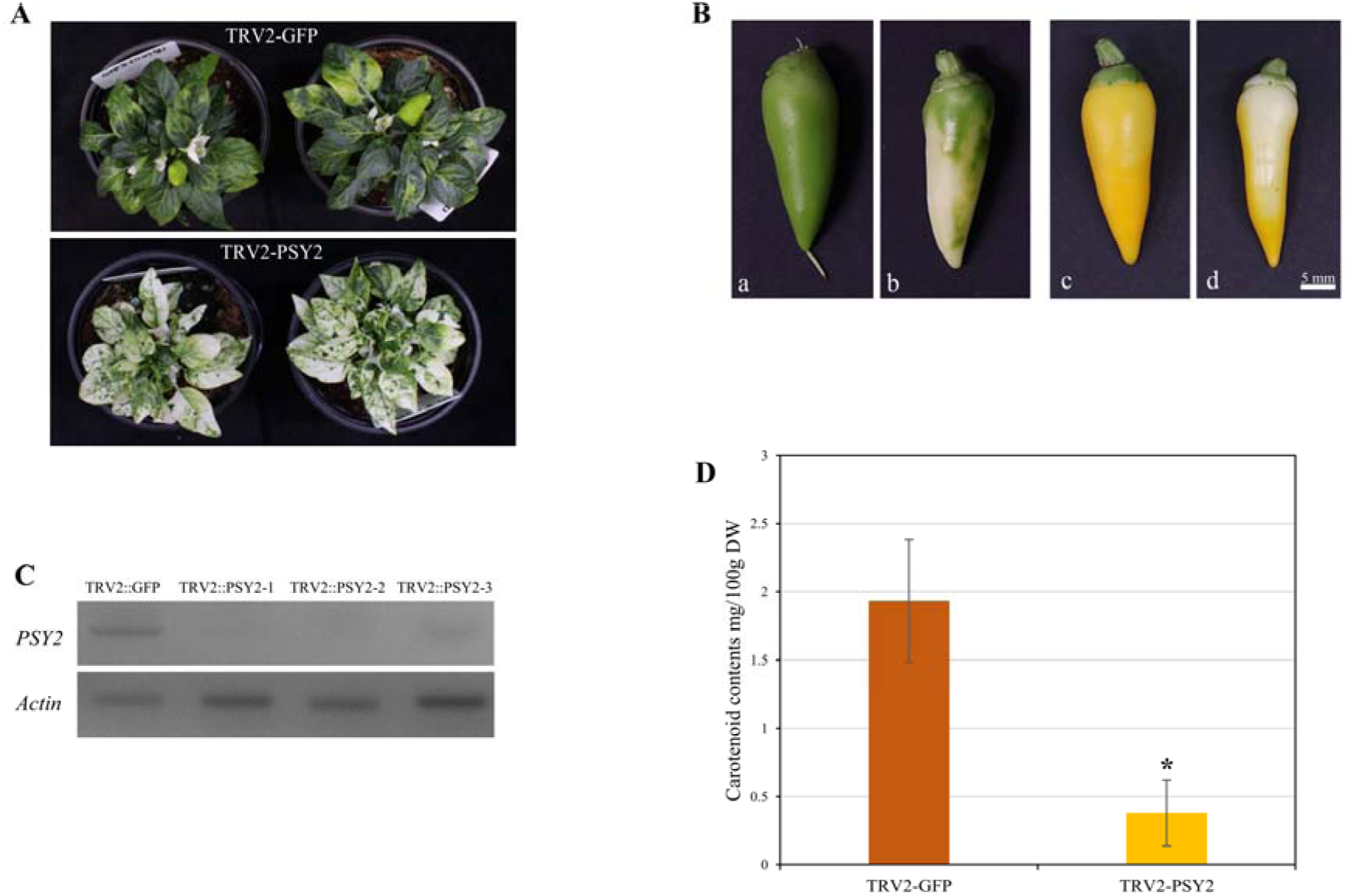
VIGS of *PSY2* in MY. (A) Phenotypes of TRV2-GFP-inoculated MY (top) and TRV2-PSY2-inoculated MY (bottom) plants at 60 days after inoculation (DAI). (B) Effect of the VIGS of *PSY2* on the color of immature (a, b) and mature (c, d) fruit. TRV2-GFP-inoculated fruits (a, c) and TRV2-PSY2-inoculated fruits (b, d). Scale bar indicates 5 mm. (C) Expression analysis of *PSY2* using RT-PCR. Three biological replicates of *PSY2*-silenced fruits were used. *Actin* serves as a control. (D) Total carotenoid contents of TRV2-GFP- or TRV2-PSY2-inoculated MY fruits. The asterisk indicates a significant difference (*P* < 0.01) from the control, determined using a Student’s *t*-test. TRV2-GFP was used as a vector control.

HPLC analysis was conducted for a quantitative and qualitative analysis of the carotenoid contents in the TRV-GFP- and TRV2-PSY2-inoculated mature fruits. The total carotenoid contents of TRV2-GFP and TRV2-PSY2 were 1.94 mg/100g DW and 0.38 mg/100g DW, respectively. The mature fruits of the control plants had a more than five-fold higher carotenoid content than the *PSY2*-silenced fruits. Lutein was detected as the most prominent component in both TRV2-GFP and TRV2-PSY2 (**Fig. 7D** and **Fig. S5**).

## Discussion

In this study, we revealed that the MY pepper line, which produces yellow mature fruits, has structural mutations in *PSY1* and *CCS*. The yellow fruit color resulted from mutation of *CCS* coding region in pepper was previously reported in various studies (Lefebvre *et al*., 1998; Popovsky and Paran 2000; Li *et al*., 2013). Our study also showed mutation in *CCS* coding region which resulted in transcriptional variant of 115 amino acids. Splicing variations and mutations in the coding sequence of *PSY1* have been analyzed previously (Kim *et al*., 2010; Jeong *et al*., 2018); by contrast, we have reported the complete deletion of *PSY1* for the first time. If PSY1 were the only enzyme able to catalyze the first step of carotenoids biosynthesis, the *psy1* knockout plants would not be able to produce carotenoids; however, we detected low levels of carotenoids in the *psy1* knockout plants. *PSY2* was expressed in the pericarp tissues of both the wild-type and *psy1* knockout plants, and we verified the enzymatic activity of PSY2 using a color complementation assay. A VIGS analysis also showed that the knockdown of *PSY2* resulted in a lighter yellow color of mature fruits of *psy1* mutant pepper. In conclusion, we showed that *PSY2* controls the yellow fruit color in pepper *psy1* knockout mutants.

Tomato has three *SlPSY* genes, *SlPSY1, SlPSY2*, and *SlPSY3* (Fantini *et al*., 2013). In contrast to *SlPSY1*, which is expressed predominantly in fruit, *SlPSY2* is expressed mainly in petals, followed by the leaf, sepal, and ovary tissues, as well as at a lower level in the remaining tissues, including the root and fruit (Giorio *et al*., 2008). *SlPSY3* is expressed mainly in root tissues under stress conditions (Kachanovsky *et al*., 2012). The silencing of *SlPSY2* and *SlPSY3* did not result in dramatic differences in the mature tomato fruit color (Fantini *et al*., 2013), suggesting that these genes have a negligible role in the pericarp tissues. In addition to *CaPSY1*, the pepper genome contains two more putative *PSY* genes, whose functions have not yet been identified. We hypothesized that *CaPSY2* might be involved in color formation in MY as *CaPSY3* was not amplified in MY fruit. Consistent with the expression pattern of *SlPSY2*, only a basal level of *CaPSY2* transcripts were obtained in the fruit, regardless of the developmental stage analyzed (**Fig. 5B**). Meanwhile, the *CaPSY2* transcript level was much higher in MY than in MR in most tissues, indicating the positive feedback and corresponding upregulation of this gene in MY due to the absence of *CaPSY1*. We revealed the enzymatic activity of *Ca*PSY2 using a color complementation assay in *E. coli.* Similar to previous studies using pAC-85b to confirm the activity of *PSY* homologs in other crops, we first investigated the color change of the cell pellet and performed an UPLC analysis of the cell extract. Both results demonstrated that *Ca*PSY2 functions as an enzyme in carotenoid biosynthesis, and we concluded that *Ca*PSY2 can supplement the activity of *Ca*PSY1 and contribute to the development of the yellow color in MY fruits.

Using single-molecule real-time (SMRT) sequencing, Jeong *et al*. (2018) identified the sequence variations of six carotenoid biosynthetic genes, including *PSY1* and *PSY2*, in diverse *Capsicum* genotypes. Of the 94 *C. annuum* accessions they analyzed, *PSY1* was not amplified from 18 lines. We hypothesized that these genotypes could have a structural variation identical to the large deletion revealed in the present study. The same deletion of *PSY1* was found in 15 out of 18 accessions that previously did not showed *PSY1* amplification using SCAR marker designed in this study (**Fig. S4**). The marker developed in this study is therefore widely applicable for use in other *C. annuum* germplasms. Moreover, Jeong *et al*. (2018) reported another three *psy1* knockout mutations in eight accessions, two of which are frame-shift mutations (*psy1-c5* and *psy1-c10*) and the other is a nonsense mutation (*psy1-c9*). There were, however, no critical mutations identified in *PSY2*. These results also support our hypothesis that *PSY2* may complement the function of *PSY1* in non-functional *PSY1* accessions.

According to the three-locus model, the diverse colors of pepper fruits are determined by *PSY1*, *CCS*, and one unidentified locus (Lefebvre *et al*., 1998; Huh *et al*., 2001). It is not possible to state that *PSY2* is encoded by the last locus based solely on the results of our study, as we have not found the natural *PSY2* mutation with mutant *PSY1*. Today, other aspects of carotenoid metabolism have been investigated for their role in fruit colors of pepper. In these studies, several transcription factors have been revealed to play a role in the color intensities of mature pepper fruits. The Golden2-like transcription factor (GLK2) and Arabidopsis pseudo response regulator 2-like (APRR2) affect plastid levels and thus regulate color in a quantitative manner in pepper fruit (Brand *et al*., 2012; Pan *et al*., 2013). To obtain clear evidence of the function of *PSY2* in fruit color formation, further experiments were required. For this reason, we used VIGS to silence *PSY2* expression in the *psy1* knockout plants, which resulted in a white mosaic phenotype in both the leaves and fruit, supporting the other results obtained in this study.

Using an UPLC analysis, we analyzed the carotenoid profiles of two pepper genotypes producing orange fruits. The orange color of the *PSY1/- ccs/ccs* fruits resulted from the moderate accumulation of diverse carotenoid components except capsanthin and capsorubin, whereas the orange color of *psy1/psy1 CCS/-* fruits was generated by the proportionally high level of red pigments (52.0% for capsanthin, 18.9% for capsorubin), although their absolute contents are considerably lower than that of the *PSY1/- CCS/-* type. In the case of *PSY1/- CCS/-* red fruits, the proportion of these red pigments accounts for 67.5% of all carotenoids, which was slightly less than that of the pigments in the *psy1/psy1 CCS/-* orange fruits. This implies that the absolute content of the carotenoid pigments is more important for the development of red fruits than the relative proportions of red and other pigments.

In conclusion, we demonstrated that a candidate gene approach combined with chemotyping via UPLC can be a useful tool for the identification of genetic factors controlling the carotenoid metabolic pathway. While the contribution of PSY2 seemed to be minimal in pepper fruit compared with that of functional PSY1, PSY2 seems to play an essential role in the *psy1* knockout individuals, as it can allow a basal level of carotenoid accumulation. *PSY2* must therefore be considered alongside *PSY1* and *CCS* as an essential gene for the successful breeding of peppers with new color and carotenoid profiles.

## Acknowledgments

This work was supported by the Korea Institute of Planning and Evaluation for Technology in Food, Agriculture, and Forestry (IPET) through the Agriculture, Food, and Rural Affairs Research Center Support Program (Vegetable Breeding Research Center, 710011-03), funded by the Ministry of Agriculture, Food, and Rural Affairs (MAFRA). This work was carried out with the support of the Cooperative Research Program for Agriculture Science & Technology Development (Project No. PJ01322901), Rural Development Administration, Republic of Korea.

## Supplementary tables

**Table S1.**
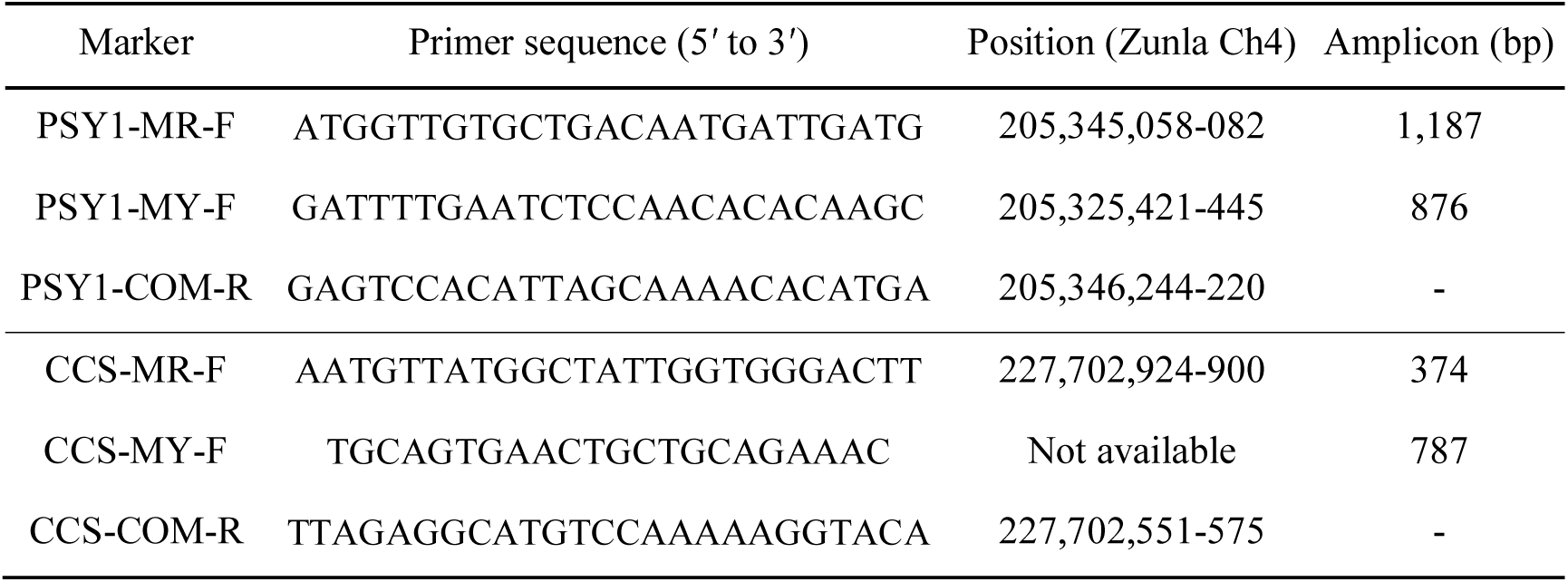
List of primers used in the SCAR marker analyses of *PSY1* and *CCS*.

## Supplementary figure legends

**Fig. S1.**
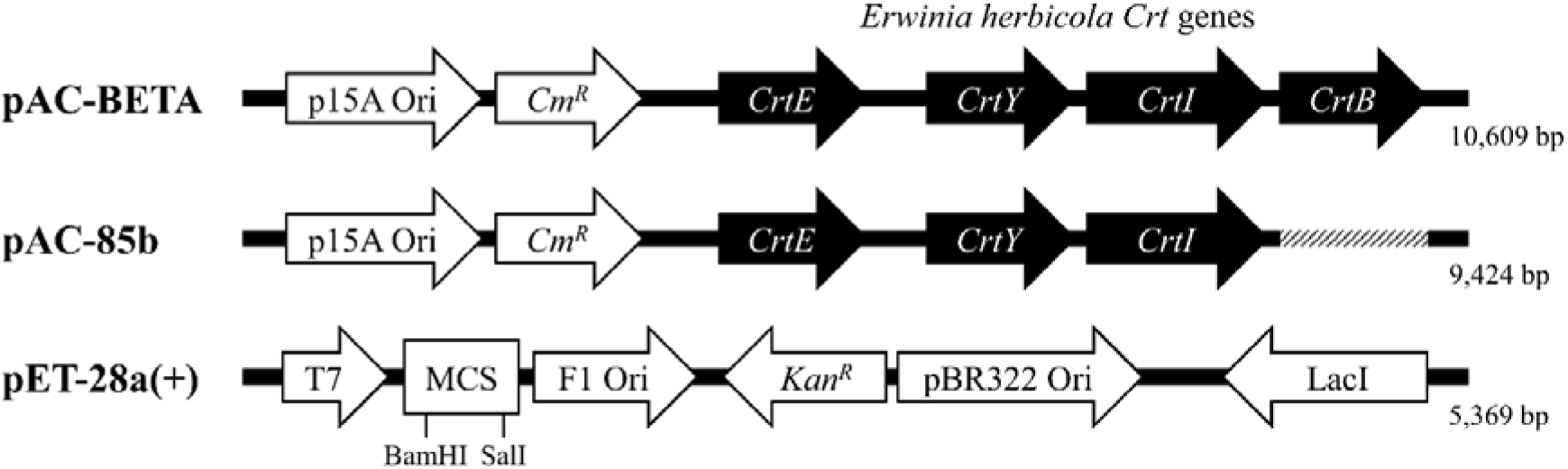
Vector information used in the color complementation assay. pAC-BETA contains the *CrtE, CrtY, CrtI*, and *CrtB* carotenoid pathway genes from *Erwinia herbicola*, and thus causes β-carotene production in *E. coli*. The pAC-85b vector lacks *CrtB,* a *PSY* homolog, and cannot cause the biosynthesis of β-carotene. For the expression of *PSY2* derived from MicroPep, the pET-28a(+) vector containing the *T7* promoter was used.

**Fig. S2.**
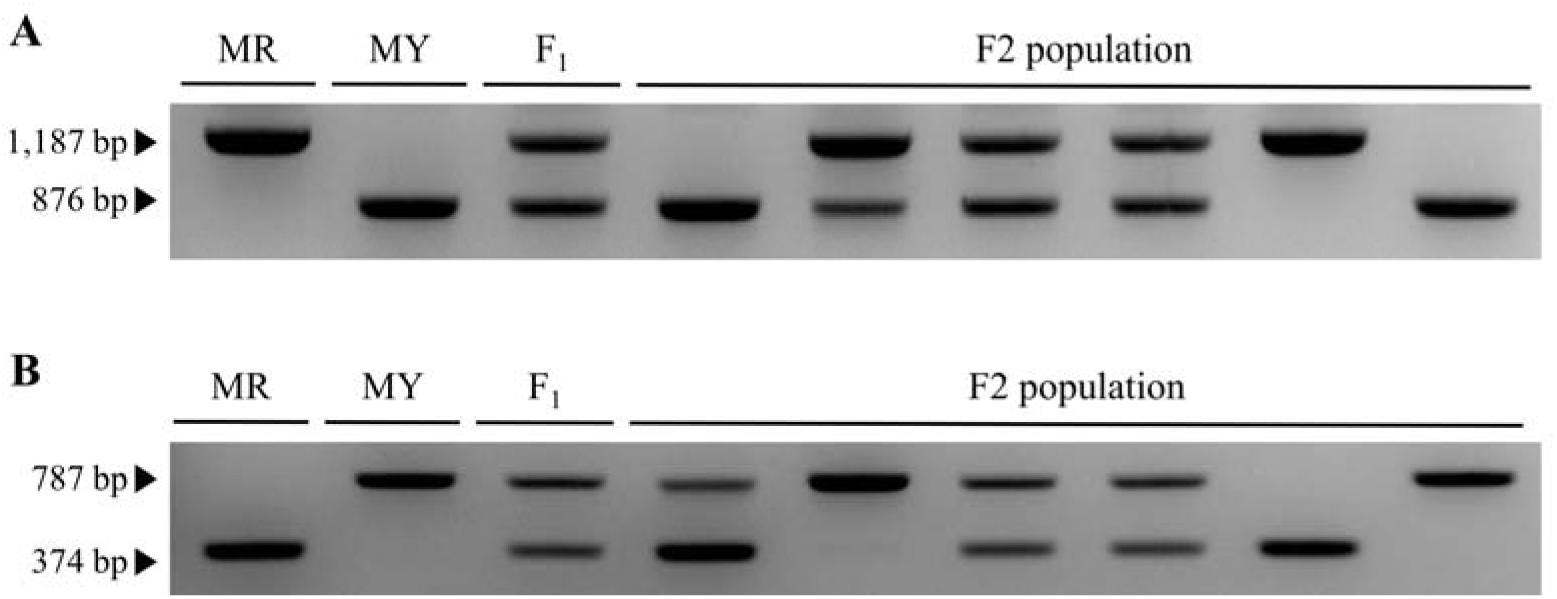
SCAR marker genotyping of *PSY1* and *CCS*. Marker genotype of *PSY1* (A) and *CCS* (B) in the parental lines MR and MY, an F_1_ hybrid, and six representative F_2_ plants. For *PSY1*, a bigger band (1,187 bp) is a representative of the MR-type allele, while a smaller band (374 bp) is a representative of the MY-type allele in *CCS*.

**Fig. S3.**
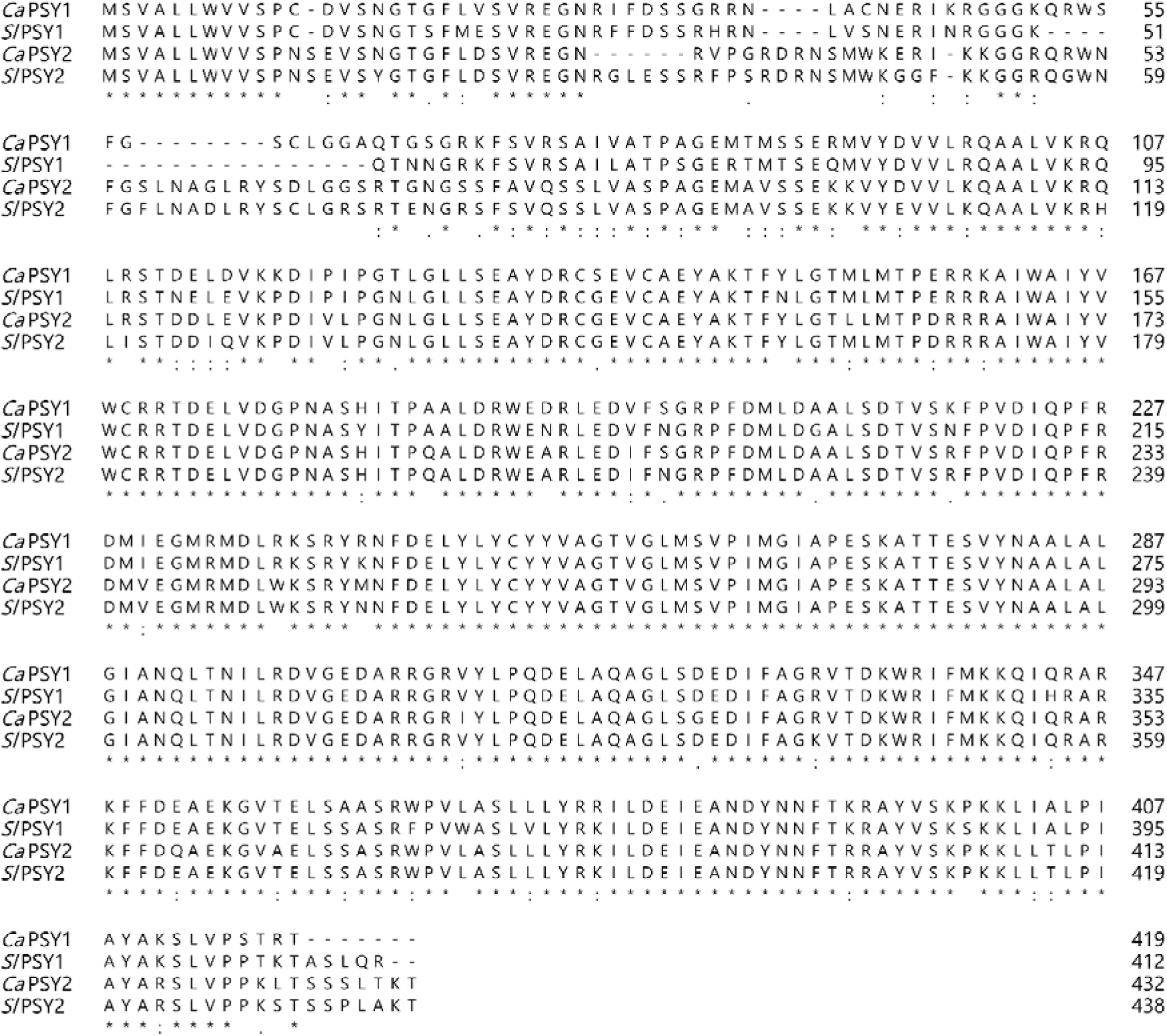
Amino acid sequence alignment of pepper and tomato PSY1 and PSY2. Asterisks (*) indicate an identical residue between the sequences.

**Fig. S4.**
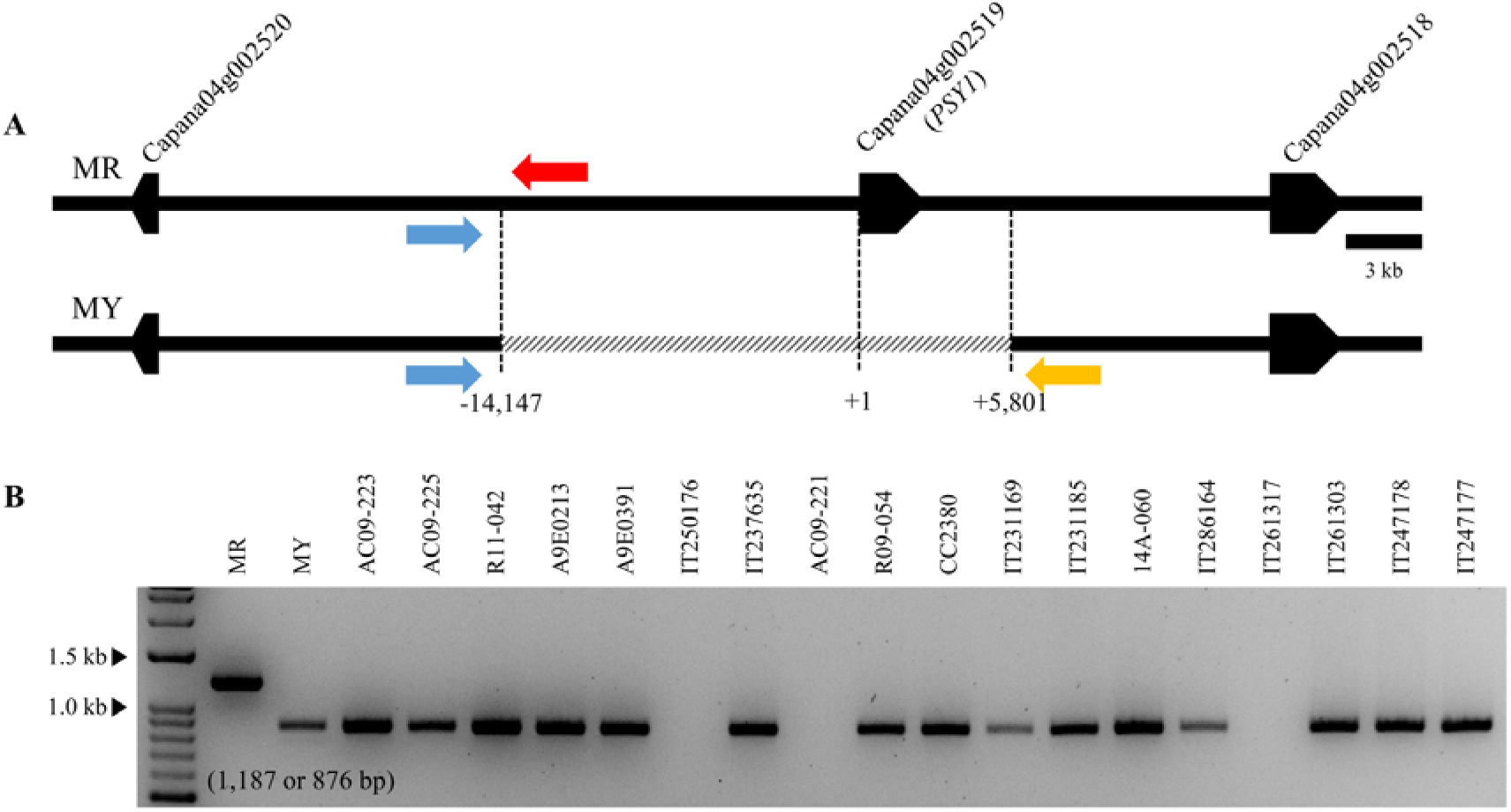
SCAR marker genotyping of the accessions from which *PSY1* was not previously amplified in study of Jeong *et al*. (2018). (A) The primer binding sites in a genomic diagram. Common forward primer (blue) and distinct reverse primers (red and yellow, respectively) were used for each template. (B) Results of the SCAR marker test. Among the 18 accessions in which *PSY1* was not amplified, 15 were revealed to possess a structural variation identical to MY. The other three accessions were not amplified.

**Fig. S5.**
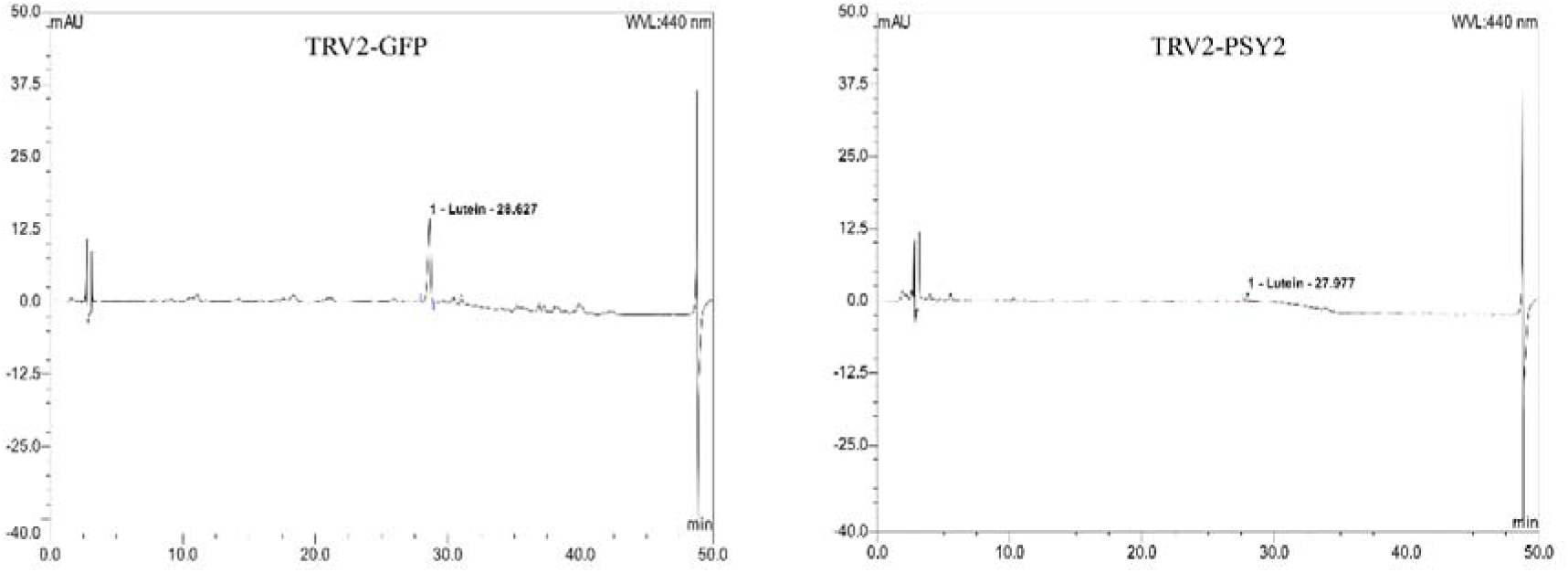
HPLC profiles of the carotenoid extracts obtained from the VIGS-treated MY plants. Both TRV2-GFP-(left) and TRV2-PSY2-inoculated (right) mature fruits contained lutein as a major component. Au, absorbance units.

**Appendix 1.**
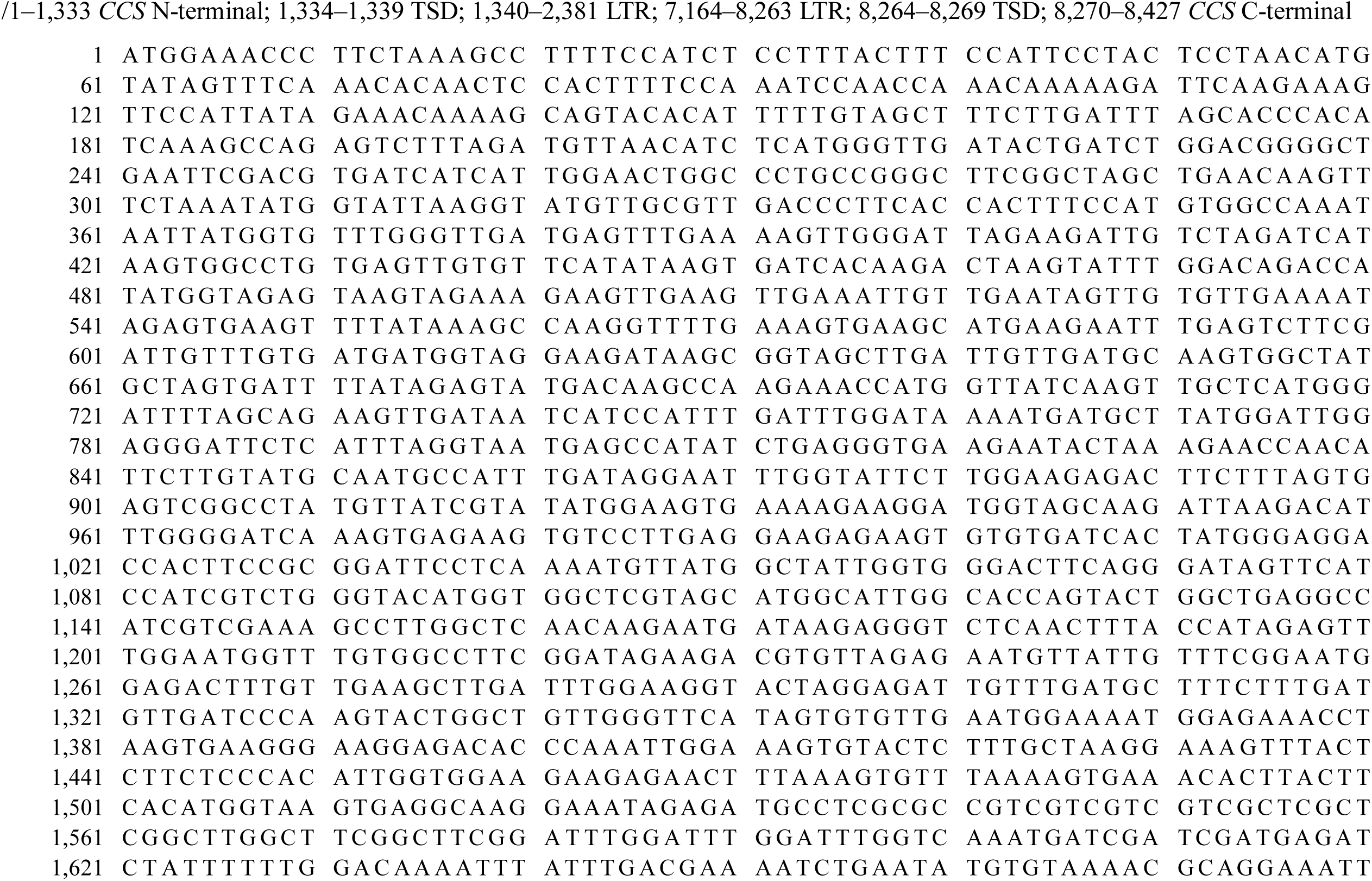
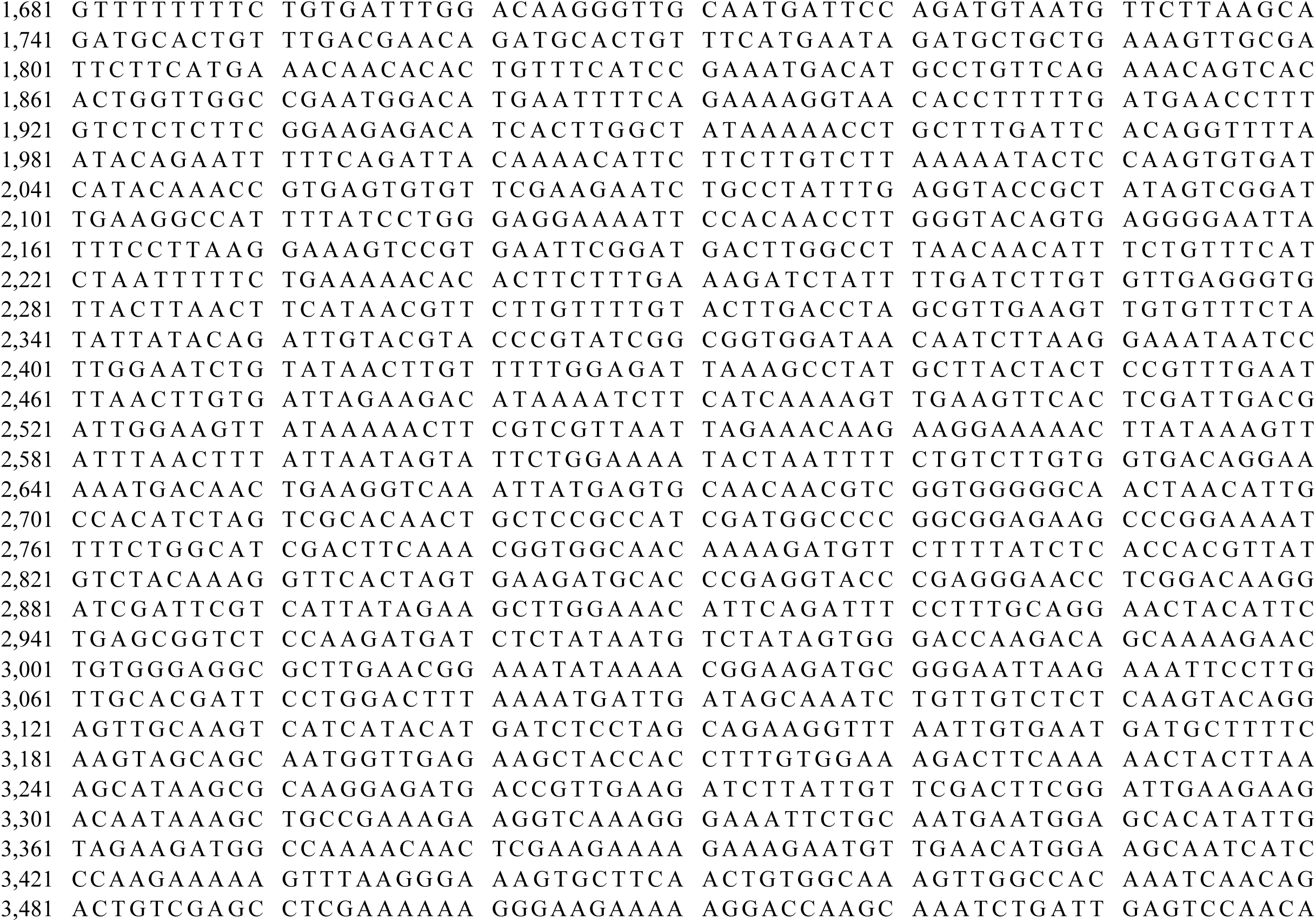
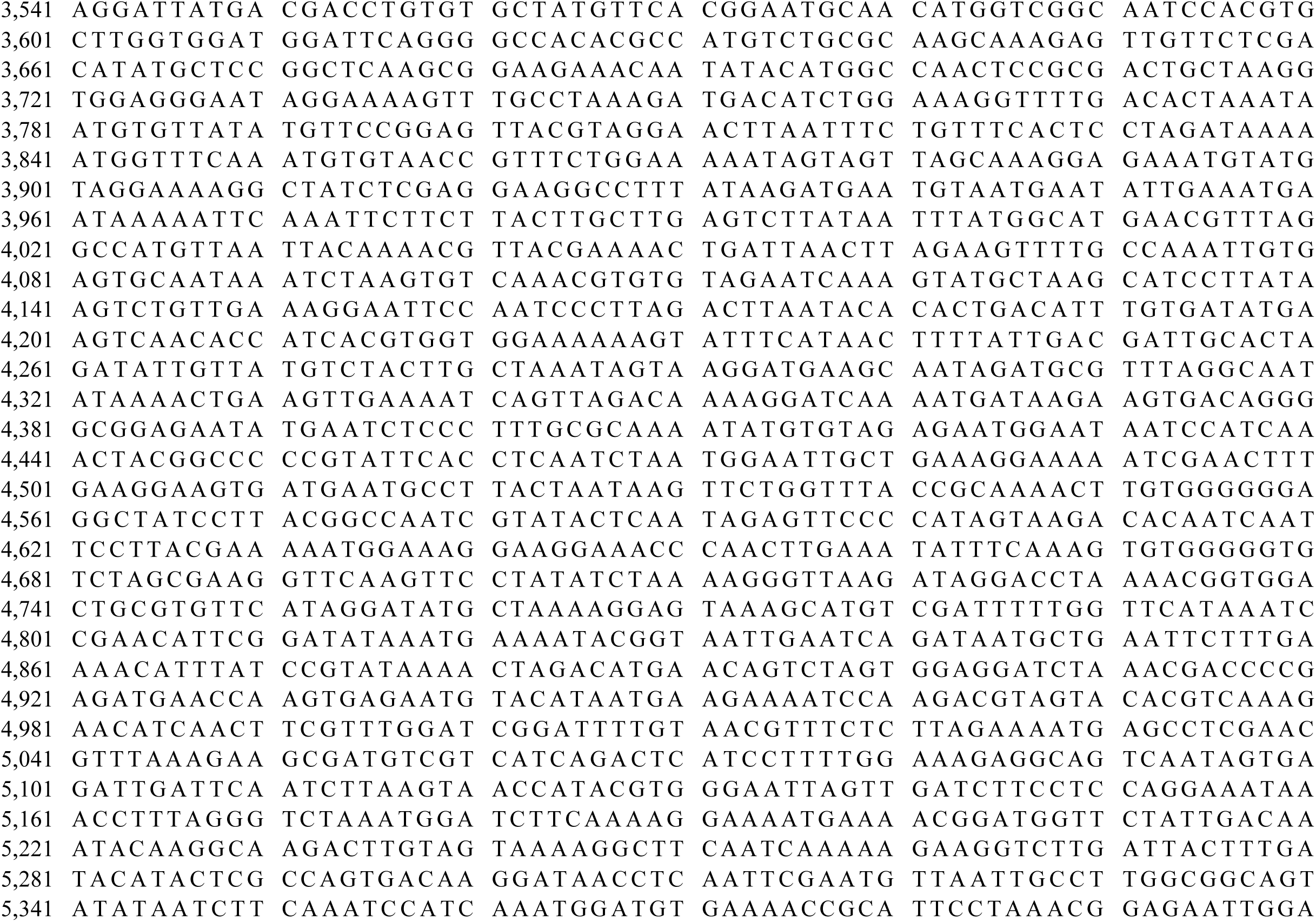
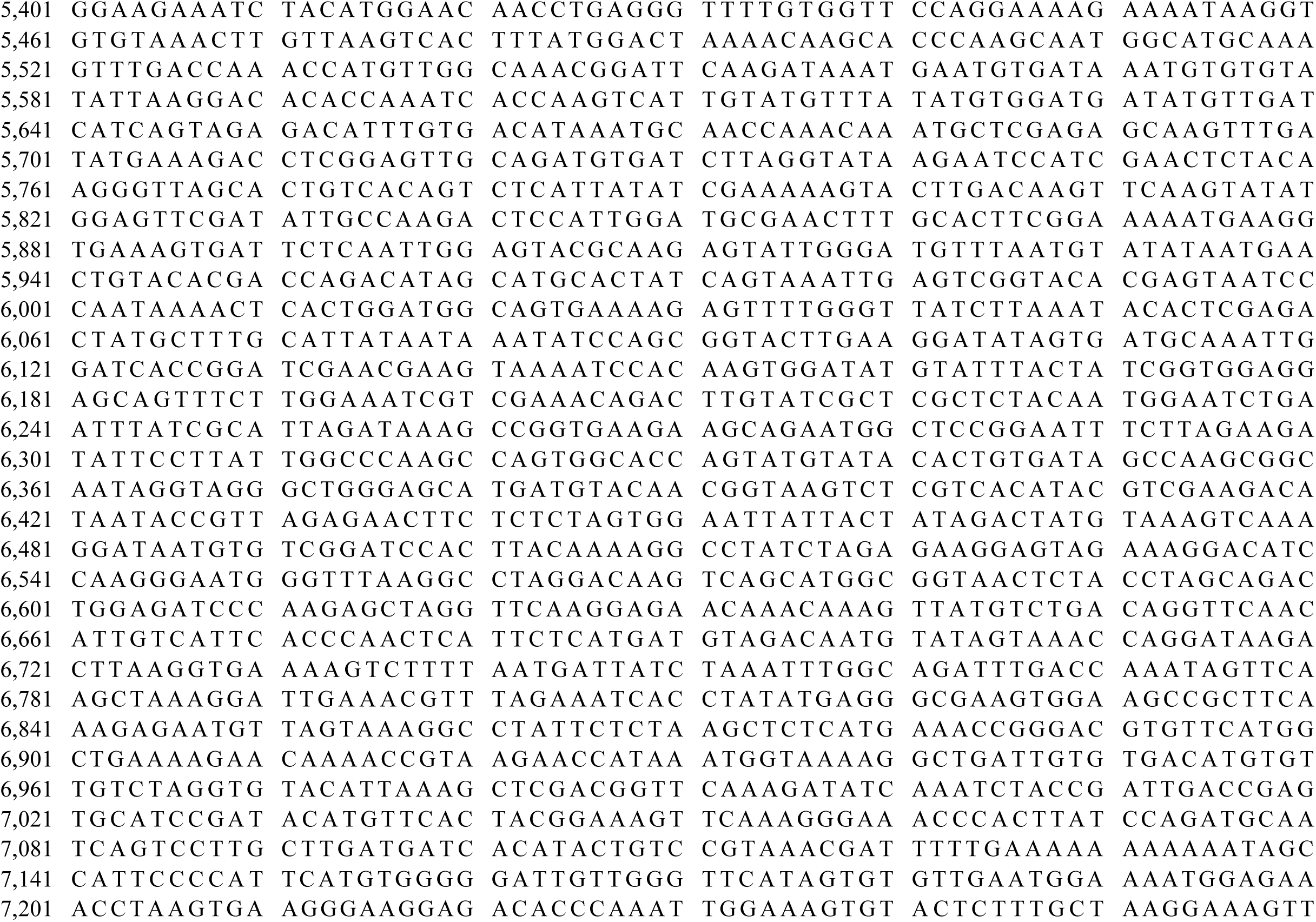
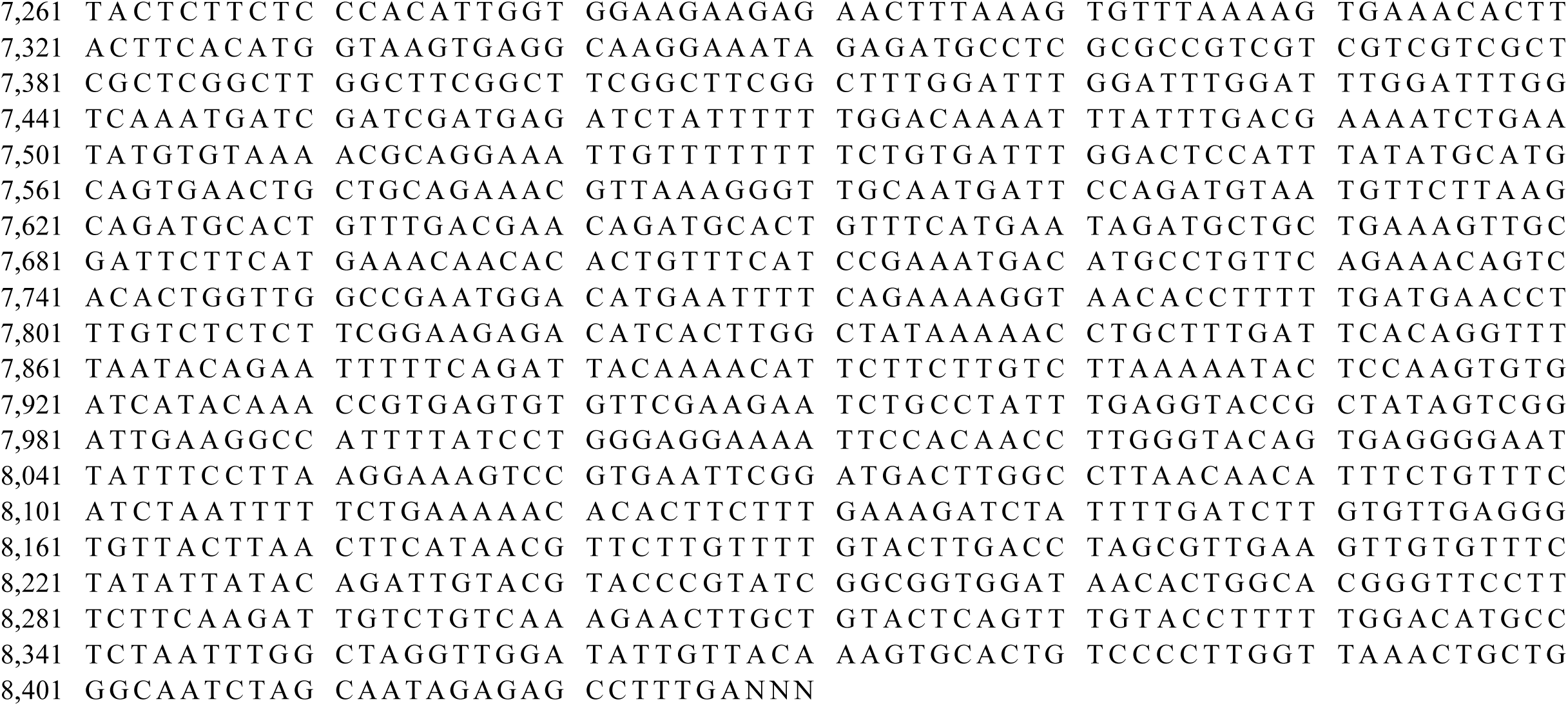
LTR retrotransposon sequences discovered in the MY CCS sequence.

